# Integrative Epigenomic Landscape of Alzheimer’s Disease Brains Reveals Oligodendrocyte Molecular Perturbations Associated with Tau

**DOI:** 10.1101/2025.02.12.637140

**Authors:** Stephanie R. Oatman, Joseph S. Reddy, Amin Atashgaran, Xue Wang, Yuhao Min, Zachary Quicksall, Floor Vanelderen, Minerva M. Carrasquillo, Chia-Chen Liu, Yu Yamazaki, Thuy T. Nguyen, Michael Heckman, Na Zhao, Michael DeTure, Melissa E. Murray, Guojun Bu, Takahisa Kanekiyo, Dennis W. Dickson, Mariet Allen, Nilüfer Ertekin-Taner

**Affiliations:** Mayo Clinic, Department of Neuroscience, Jacksonville, FL USA; Mayo Clinic, Department of Quantitative Health Sciences, Jacksonville, FL USA; Mayo Clinic, Department of Neurology, Jacksonville, FL USA

## Abstract

Alzheimer’s disease (AD) brains are characterized by neuropathologic and biochemical changes that are highly variable across individuals. Capturing epigenetic factors that associate with this variability can reveal novel biological insights into AD pathophysiology. We conducted an epigenome-wide association study of DNA methylation (DNAm) in 472 AD brains with neuropathologic measures (Braak stage, Thal phase, and cerebral amyloid angiopathy score) and brain biochemical levels of five proteins (APOE, amyloid-β (Aβ)40, Aβ42, tau, and p-tau) core to AD pathogenesis. Using a novel regional methylation (rCpGm) approach, we identified 5,478 significant associations, 99.7% of which were with brain tau biochemical measures. Of the tau-associated rCpGms, 93 had concordant associations in external datasets comprising 1,337 brain samples. Integrative transcriptome-methylome analyses uncovered 535 significant gene expression associations for these 93 rCpGms. Genes with concurrent transcriptome-methylome perturbations were enriched in oligodendrocyte marker genes, including known AD risk genes such as *BIN1*, myelination genes *MYRF, MBP* and *MAG* previously implicated in AD, as well as novel genes like *LDB3*. We further annotated the top oligodendrocyte genes in an additional 6 brain single cell and 2 bulk transcriptome datasets from AD and two other tauopathies, Pick’s disease and progressive supranuclear palsy (PSP). Our findings support consistent rCpGm and gene expression associations with these tauopathies and tau-related phenotypes in both bulk brain tissue and oligodendrocyte clusters. In summary, we uncover the integrative epigenomic landscape of AD and demonstrate tau-related oligodendrocyte gene perturbations as a common potential pathomechanism across different tauopathies.

## Introduction

Amyloid beta (Aβ) plaques and tau neurofibrillary tangles (NFT) are core neuropathological hallmarks of Alzheimer’s Disease (AD)^1,2^. The characteristic distribution and levels of these amyloid and tau hallmarks in the brain are often measured by Thal phase^3^ and Braak stage^4^, respectively. These core measures show high inter- and intra-individual brain variability within AD^5–8^ suggesting the presence of distinct mechanisms controlling the levels and biochemical states of their underlying proteins. This is supported by our previous findings that key proteins involved in the pathophysiology of AD including Aβ, tau, and apolipoprotein E (APOE) each have unique genetic architectures that associate with their specific biochemical states and levels in the AD brain^9^. Characterization of these endophenotype specific molecular mechanisms in the AD brain are important as each protein and state has potentially unique functions and consequences^10^. Insoluble states of Aβ42 and Aβ40 are major components enriched in Aβ plaques in the brain parenchyma and cerebral amyloid angiopathy (CAA) deposits, respectively^6,11^. CAA is a co-pathology often seen in AD characterized by amyloid deposits in the cerebral vasculature which can impair blood flow and may contribute to clinical deficits beyond AD^12–14^. Insoluble states of tau, particularly hyperphosphorylated tau (pTau), are core components of NFT which are often seen after brain levels of soluble total tau rise and become hyperphosphorylated^15^. Recent studies have also shown that membrane bound states of Aβ and tau correlate with clinical severity and that tau increases its propensity to fibrillize when contacting a plasma membrane^16–21^. Moreover, the APOE protein which is important in transporting and processing lipids in the brain, is co-deposited in insoluble Aβ plaques and its gene harbors the largest known genetic risk factor for AD, the *APOEε*4 tagging variant^22–24^.

Although there has been great progress identifying risk for AD including genetic and environmental/lifestyle factors^25,26^, the underlying molecular mechanisms that these risk factors act through to impact specific AD endophenotypes remains to be fully characterized. It is likely that these risk factors affect both shared and distinct mechanisms that underpin the hallmarks of AD and investigating these factors will be important in discovering the molecular underpinnings of AD. Epigenetic mechanisms are known to mediate genetic as well as environmental and lifestyle factor effects on gene expression. Previous epigenetic studies of AD found broad dysregulation of epigenetic mechanisms corresponding to transcriptomic dysregulation^27,28^. A key epigenetic regulator of gene expression is DNA methylation (DNAm). Previous studies have found differential DNAm sites near known AD risk genes including *BIN1, APOE, APP, ADAM10, MAPT, SPI1, TREM2* and others^29–37^. DNAm changes have also been linked to AD risk, onset and progression^30–32,38,39^. These results suggest that epigenetic regulation, particularly that of DNAm, may be important in modulating expression of known AD risk genes and thereby influencing core AD endophenotypes. Nonetheless, large scale, integrative studies have yet to investigate how dysregulation of DNAm impacts specific brain endophenotypes beyond global neuropathology levels. Identification of AD-endophenotype specific molecular perturbations is required to deconstruct the underlying pathophysiology of AD.

We hypothesize that, within the AD brain, neuropathology, levels and biochemical states of core AD-related proteins are influenced by variations in DNAm. To test this, we performed epigenome-wide association studies (EWAS) using an innovate approach to analyze regional levels of DNAm (rCpGm) from the temporal cortex (TCX) and cerebellum (CER) with AD-related neuropathologic measures (Braak, Thal, and CAA) and brain biochemical levels of five proteins (APOE, Aβ40, Aβ42, tau, and pTau) (**Figure 1**). Through our deep endophenotype EWAS and integrative omics approach leveraging a large cohort of neuropathologically confirmed AD cases, we identified widespread associations of DNAm with levels of distinct biochemical states of tau connected to significant dysregulation of nearby oligodendrocyte and myelin related gene expression. Interrogation of these tau related, epigenetically dysregulated oligodendrocyte genes and loci revealed concordant tau related associations across multiple independent datasets of AD and other tauopathies. Our findings suggest a key relationship between brain levels and biochemical states of tau and expression of oligodendrocyte genes regulated through DNAm perturbations. This may be a common pathomechanism across AD and other tauopathies.

**Figure 1:**
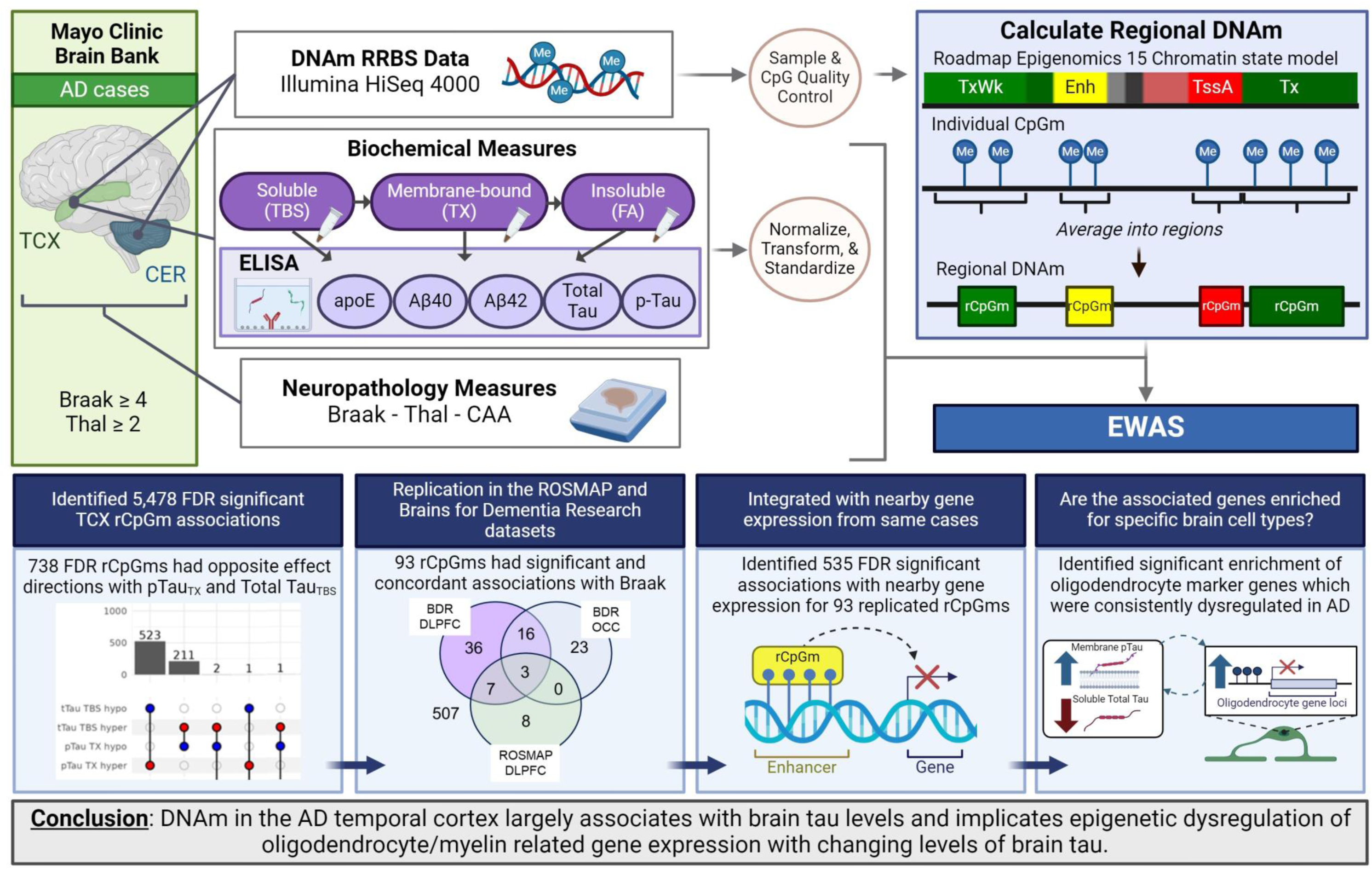
Graphical abstract. Graphical description of the study design. Neuropathologically diagnosed AD cases were identified from the Mayo Clinic Brain Bank. Samples had neuropathology measures, biochemical measures from the temporal cortex, and DNA methylation data from the temporal cortex and a subset of samples from the cerebellum. The DNA methylation data was grouped into regions based on windows defined by the Roadmap Epigenomics 15 Chromatin state model temporal lobe predictions (Kundaje *et al.* 2015). These rCpGms were tested for association with the biochemical and neuropathology measures. Top EWAS hits were then investigated identifying important perturbed biological pathways related to tau and oligodendrocyte gene expression. **AD**= Alzheimer’s Disease; **CAA**= Cerebral Amyloid Angiopathy; **CpGm** = DNA methylation at CpG sites; **Enh** = Enhancer; **EWAS**= Epigenome wide association study; **FA**= Formic Acid; **Me** = Methylation; **rCpGm** = Regional DNA methylation grouped by chromatin state; **RRBS** = Reduced Representation Bisulfite Sequencing; **SNP** = Single Nucleotide Polymorphism; **TBS**= Tris Buffered Saline; **TssA**= Transcription Start Site Active; **TX**= Triton-X; **Tx** = Strong Transcription; **TxWk** = Weak Transcription; Adapted from Oatman & Reddy et al. 2023, *Molecular Neurodegeneration*; Created in Biorender.com.

## Results

We analyzed levels of DNA methylation at CpG sites (CpGm) using reduced representation bisulfite sequencing (RRBS) from two brain regions, TCX and CER, in a dataset of neuropathologically confirmed AD donors with multiple AD-related endophenotype measurements. These endophenotypes included neuropathological variables (Braak stage, Thal phase, and CAA scores) and TCX biochemical measures of five AD-related proteins^10^ (APOE, Aβ40, Aβ42, total tau, and phospho-Tau (pTau) (Thr231)) from three tissue fractions (soluble (TBS), membrane-bound (TX), and insoluble (FA)) (**Figure S1**). Our study dataset consisted of 472 individuals (53% female) with CpGm measured from the TCX and a subset of 200 from the CER (**Table S1**). After stringent quality control and data reduction steps, a total of 455 TCX and 191 CER samples with 1,958,174 TCX and 1,858,631 CER CpGm sites were available for downstream analytics (**Table S2, Figure S2**).

### Grouping CpGm by predicted chromatin state is a biologically relevant method to investigate regional levels of DNA methylation

A commonly used EWAS approach to identify AD-related CpGm loci is to associate individual CpGm sites with a phenotype of interest. However, this individual CpGm based EWAS does not account for methylation variations at neighboring CpGm sites which are highly correlated and likely work together to have a functional effect^40–43^. Currently available methods address this by grouping nearby significant individual CpGm sites into regions, however, these approaches are based on post-hoc statistical groupings which lack biological basis^44–47^. In this study, we aimed to investigate CpGm on a regional level defined by the biological function of the genomic region to identify functionally relevant CpGm loci. To do this, we developed a novel method that averages CpGm sites into regions (rCpGm) based on biologically based genomic windows defined by the Roadmap Epigenomics Consortium 15-chromatin state model generated from the human temporal lobe^48^, a region pertinent to our human brain dataset (**Figure 1**).

First, to determine whether our novel method yields biologically consistent results, we investigated the average methylation levels of rCpGm sites grouped by brain chromatin states^48^. We found that the average methylation levels for each chromatin state were highly consistent with the predicted biology of these states (**Figure 2**). Quiescent, repressed, and heterochromatin annotations had higher methylation, typically associated with lower gene expression levels^49^, whereas active transcription start sites (TSS), flanking TSS, and bivalent/poised TSS regions had low methylation levels. Chromatin regions that are expected to have more variability across cell types, like enhancer regions, have around 50% methylation in our bulk brain tissue rCpGm data. Our chromatin state specific rCpGm averages were also highly consistent with published whole-genome bisulfite sequencing (WGBS) DNA methylation data^48^ demonstrating the utility of our method for defining biologically relevant CpGm regions.

**Figure 2:**
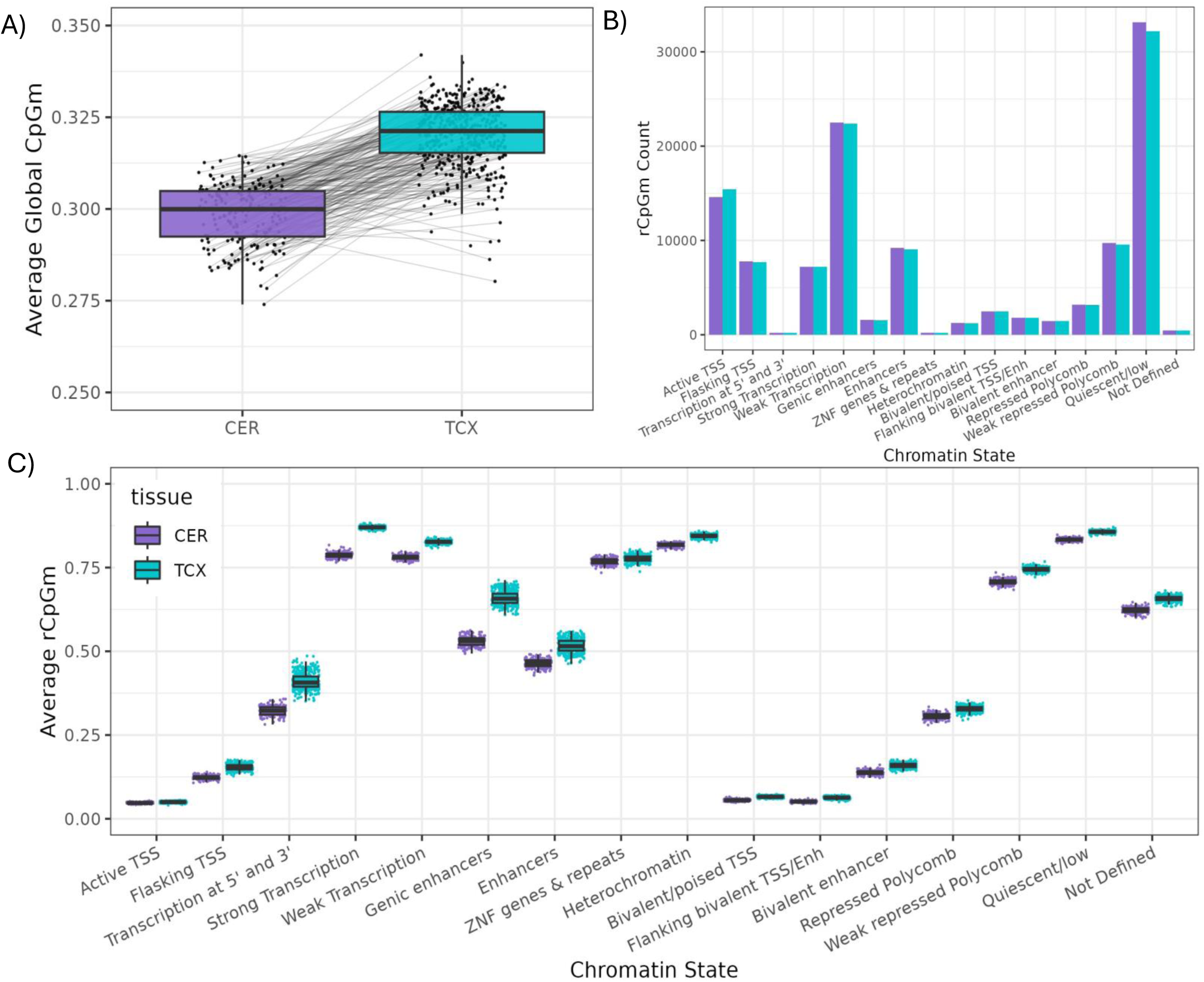
Characterization of global DNAm and identified rCpGms in our dataset. A) Average global CpG methylation calculated for each sample before data reduction QC steps. Samples with both CER and TCX CpGm measures (N= 190) are connected by a grey line. B) Number of rCpGm sites in the CER and TCX that were annotated to each predicted chromatin state. C) Average rCpGm for each sample split by chromatin state and tissue type. Chromatin states annotated as “Not Defined” are the regions that did not lift over from hg19 to hg38 in the Epigenomics Roadmap model and so do not have a defined annotation.; Using two sample t-test, we found the TCX and CER average rCpGm across each annotation were also significantly different from each other (pvalue < 1E-05) as well as the average global CpGm between the CER and TCX. **CER =** Cerebellum; **Enh**= Enhancer; **rCpGm** = CpG methylation grouped by predicted chromatin state; **TCX** = Temporal Cortex; **TSS**= Transcription Start Sites

Next, we compared the average rCpGm levels between the TCX and CER brain regions for each chromatin state annotation. TCX and CER have mostly similar levels of rCpGm for each chromatin state with the largest differences in enhancer and transcription states which may reflect the distinct cell-specific epigenetic regulation of these two brain regions (**Figure 2**). Investigating the 190 donors with both TCX and CER rCpGm measures, we found that average global rCpGm levels for each sample were similar between these brain regions (range: 0.27-0.34) and highly correlated in the same individual (r = 0.88 – 0.94, p-value < 2.2e-16), although TCX had consistently higher global methylation levels than CER (**Figure 2**, **Figure S2**).

We next explored the applicability of our novel method in two independent brain DNAm datasets measured via array technologies which are more often used in large-scale AD EWAS. These two datasets are the ROSMAP^30^ (n=686) and Brains for Dementia Research (BDR)^32^ cohorts with CpGm measured from the dorsolateral prefrontal cortex (DLPFC) in both datasets and also the occipital cortex (OCC) in BDR (n_DLPFC_=581, n_OCC_=576) (**Table S3**). Quality control (QC) processing of the DNAm levels from these datasets through our array-specific pipeline resulted in 429,723 CpGm sites in ROSMAP and 747,577 CpGm sites in BDR. Using our novel rCpGm method, we identified 136,201 rCpGms in ROSMAP and 203,435 rCpGms in BDR (**Figure S3**). These independent datasets are strikingly similar in the levels and patterns of their average rCpGm for each chromatin state annotation across these datasets (**Figure S3**) and in comparison to our dataset (**Figure 2**), despite differences in donors, brain regions, collection sites and methylation measurement methods highlighting the replicability of our novel analytic approach. Notably, the array-based rCpGms have a narrower range of values (**Figure S3**) in comparison to our RRBS-based (**Figure 2**) measurements, likely due to lower precision and dynamic range for the former methodology^50,51^. Further, rCpGm levels for each chromatin annotation are more similar between DLPFC and OCC regions for the BDR cohort than for the DLPFC region across BDR and ROSMAP cohorts (**Figure S3**). This suggests that inter-individual differences may be greater than intra-individual brain region differences in methylation.

Collectively, these results underscore the robust and replicable rCpGm measurements obtained with our analytic approach which demonstrate the broad utility and biological relevance of our novel method.

### DNA methylation associates primarily with tau related brain endophenotypes

To determine DNA methylation (DNAm) associations with AD-related endophenotypes we took two EWAS approaches, namely the more common method of individual CpGm associations and regional methylation associations using our novel method that averages CpGms into regions based on the 15-chromatin state model^48^.

Neuropathology endophenotypes (Braak stage, Thal phase, CAA score) were available for all study donors (i.e. both TCX and CER samples), whereas brain biochemical endophenotypes (protein levels of APOE, Aβ40, Aβ42, Total tau, and pTau measured in soluble (TBS), membrane-bound (TX), and insoluble (FA) brain tissue fractions) were measured only from TCX tissue (**Figure S1**). We performed EWAS of individual CpGm levels from both TCX and CER with neuropathology and those from TCX also with biochemical endophenotypes adjusting for age at death, sex, and the first 3 genetic principal components. We identified 2,489 FDR significant TCX CpGm associations with 10 endophenotypes and 50 FDR significant CER CpGm associations only for Thal (**Figure S4, Table S4, Table S5**). The majority (97%) of the FDR significant TCX CpGm associations were with tau biochemical measures (n=2,415). The most frequently associated endophenotype was Total tau_TBS_ with 2,268 FDR significant TCX CpGms, 21 of which were also associated with pTau_TX_. The remaining FDR significant CpGms were uniquely associated with a single endophenotype. Top TCX CpGm associations for each endophenotype were located in or near the genes *ACSF3* (Total tau_FA_), *SLC16A1-AS1* (Total tau_TBS_), *YES1* (AB40_TX_), *TRAPPC14* (Thal phase), *HR* (pTau_TX_), *RGS10* (APOE_TX_), *PLEKHJ1* (AB42_TX_), *CIB1/GDPGP1* (APOE_FA_), *ARPC1B* (AB42_FA_), and *UMODL1* (APOE_TB_)_S_. Interestingly, CpGms near *ACSF3, SLC16A1, HR, RGS10*, *PLEKHJ1*, *ARPC1B* and *UMODL1* have previously been identified in meta-analyses with Braak Stage, although for different CpGm positions^32,52^. *ACSF3, ARPC1B, CIB1, HR, PLEKHJ1, RSG10, UMODL1* also had nominal CpGm associations with CERAD score and Thal phase^32^. Our top 5 CER CpGm associations with Thal phase were near genes *ANKRD33*, *LINC02314, RAB30, ASIC1,* and *UBE2Q2*. A previous meta-analyses of cortical tissue found CpGms near *RAB30* and *UBE2Q2* to be nominally associated with Braak stage and Thal phase, respectively^32^.

Using our novel regional methylation analysis method, we performed EWAS with 116,014 TCX and 116,811 CER rCpGms with the same endophenotypes. We identified 5,478 FDR significant TCX rCpGm associations for 5 endophenotypes (Total tau_TBS_, pTau_TX_, pTau_FA_, APOE_TX_, Aβ42_FA_) (**Figure 3**, **Table S6**). There were no FDR significant TCX or CER rCpGm associations with any of the neuropathology endophenotypes **(Table S6, Table S7)**. Concordant with the individual CpGm findings, the majority of the significant TCX rCpGm associations were with Total tau_TBS_ (n=4,424) followed by pTau_TX_ (n=1,029). There were 738 rCpGms significantly associated with brain levels of both Total tau_TBS_ and pTau_TX_. All 738 rCpGms had opposite directions of association with Total tau_TBS_ and pTau_TX_, which is consistent with the negative correlation of these two endophenotypes (ρ= −0.21, p-value <0.001)^10^. This suggests that Total tau_TBS_ and pTau_TX_ may have shared but opposing molecular mechanisms that impact their levels in the brain. Each FDR significant rCpGm was on average composed of 23 individual CpGm sites (range: 1 - 895 CpGms) indicating that most of these associations are not driven by a single CpGm but are a composite methylation measure of the region. The majority (69%) of the FDR significant TCX rCpGm associations reside in more active chromatin regions including active TSS (TssA, N = 578), Flanking TssA (N = 652), transcription at gene 5’ and 3’ ends (N = 47), Strong transcription (N = 175), weak transcription (N = 707), genic enhancer (N = 256) and enhancer (N = 1363) annotations (**Figure 3**) suggesting that these significant rCpGms may be functionally important in gene expression regulation.

**Figure 3:**
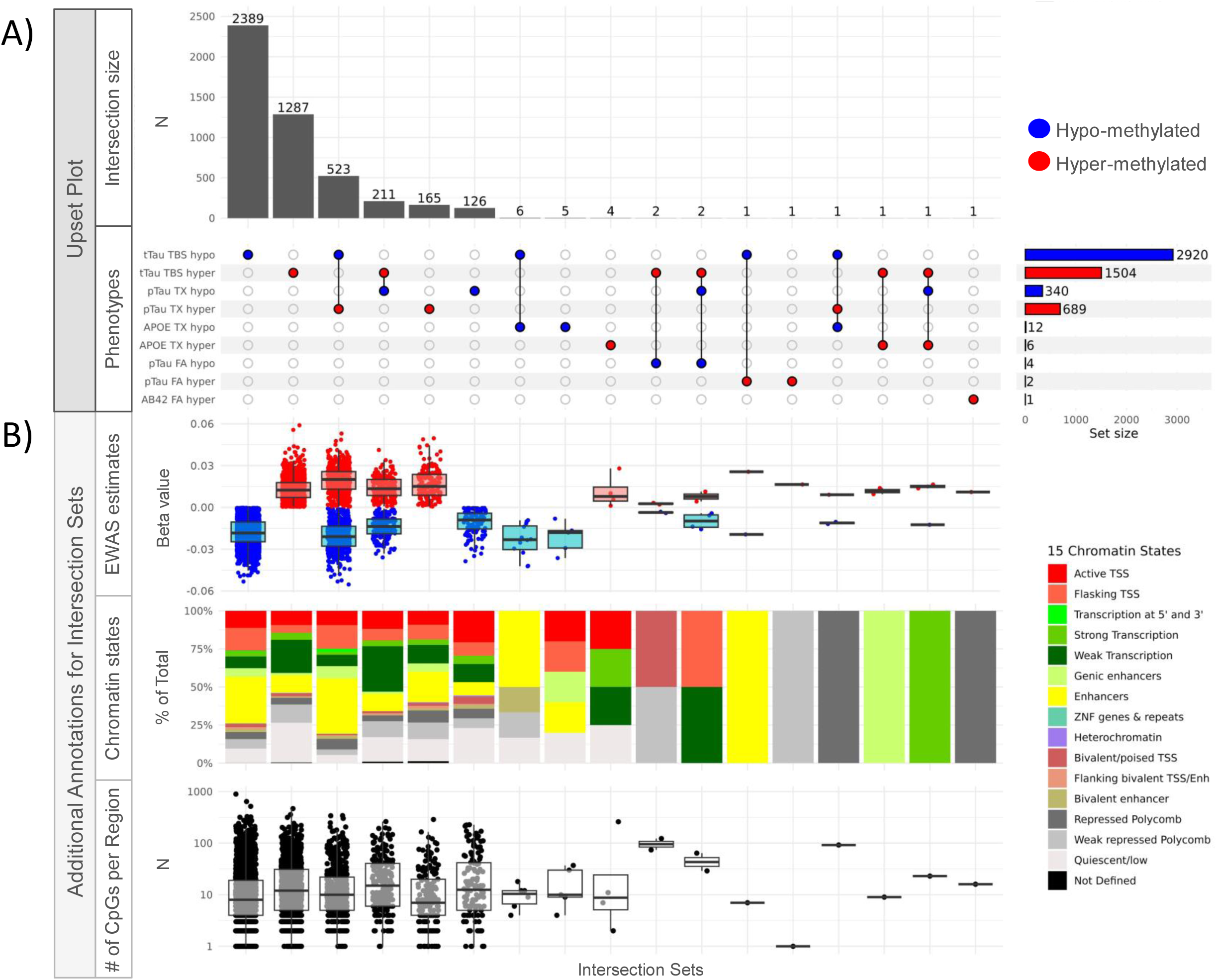
FDR significant rCpGm associations in the temporal cortex across AD endophenotypes. A) Upset plot showing all FDR significant rCpGm associations across the AD endophenotypes tested in the temporal cortex. B) Boxplot and bar graphs showing additional annotations for each intersection set of the FDR significant associations. EWAS estimates show the regression beta value for each FDR significant association colored by direction of effect. Chromatin states depict the Epigenetic Roadmap 15-chromatin state model annotation breakdown for each intersection set. ‘Number of CpGs per region’ depicts the number of individual CpGms that make up the rCpGm tested. Red indicates hyper-methylation or a positive direction of association and blue indicates hypo-methylation or a negative direction of association with the tested endophenotype. **FA** = Insoluble formic acid fraction; **FDR** = false discovery rate; **N** = Number; **pTau** = phospho-tau; **TBS** = soluble tris buffered saline fraction; **TSS** = Transcription start site; **tTau** = total tau; **TX**= membrane associated Triton-X fraction

To ensure that these FDR significant rCpGm associations are robust to any biological and technical variations, we repeated these analyses after further adjustments for Braak stage and Thal phase, neuronal cell type proportion estimates, and sequencing batch. Comparisons between the primary and further adjusted models by Pearson correlation revealed high, significant positive correlations of both the coefficient estimates (r = 0.998-0.9997) and p-values (r = 0.66-0.989) (**Figure S5**). We also investigated if the normal distribution assumptions were violated or if these associations were driven by outliers via a non-parametric Spearman rank correlation test. We found 99.3% (5,442/5,478) of the FDR significant associations also had a nominally significant (p-value ≤ 0.05) association in the Spearman rank test (**Figure S5**). These highly consistent test statistics between our primary and adjusted models and their persistent significance in a non-parametric test demonstrate the robustness of these associations to potential biological and technical confounders.

Although both the individual CpGm and our novel rCpGm analyses found the most associations with brain tau biochemical endophenotypes, there were twice as many FDR significant loci using the latter approach. Of the 2,489 FDR significant individual TCX CpGms, 68% were included in at least one of the FDR significant rCpGms. Notably, 1,671 individually significant CpGms were part of the 1,000 rCpGms associated with Total tau_TBS_, and 26 such CpGms were within 23 rCpGm associated with pTau_TX_. Thus, our novel rCpGm approach identifies both those loci captured by the more traditional single CpGms but importantly also many others that are missed by the latter. This suggests that clustering CpGms based on chromatin states is both a more powerful EWAS approach and also provides biological context for these associations. Consequently, we focused our subsequent analyses on the rCpGm results and approach. Importantly, regardless of the CpGm analytic approach, most DNAm across the genome primarily associate with tau protein levels in the AD brain.

### Replication of DNAm associations with tau neuropathology and other AD-related endophenotypes in independent datasets

We next sought replication of our findings in the ROSMAP^30^ (n=686) and Brains for Dementia Research (BDR)^32^ (n_DLPFC_=581, n_OCC_=576) cohorts that have DLPFC DNAm measured with the Illumina 450K Human Methylation and both DLPFC and OCC DNAm measured with the EPICv1 850K arrays, respectively (**Table S3**). Although to our knowledge there are no DNAm datasets that also have the precise brain biochemical measures of AD-related proteins available in our dataset, ROSMAP and BDR are two of the largest cohorts with both brain DNAm and AD-related endophenotypes.

We already demonstrated the broad similarities of rCpGm levels and patterns for each chromatin state annotation across these datasets and ours (**Figure S3**, **Figure 2**), despite the fact that the individual CpGm sites measured by their DNAm platforms are not identical. To determine whether overlapping rCpGms that are obtained from different datasets correlate with each other, we performed pairwise correlations (**Figure S6**). Using DNAm levels averaged across all samples within each dataset for all rCpGms overlapping with those from the other datasets, we conducted Spearman correlations. There are strong positive correlations for DNAm levels of overlapping rCpGm sites for each pairwise dataset analysis. As expected, these correlations are strongest for BDR DLPFC and BDR OCC datasets (Spearman Coefficient=0.996) obtained from the same brain donors, followed by BDR DLPFC and ROSMAP DLPFC that have measurements from the same brain region both using similar array technologies (Spearman Coefficient=0.914). Nonetheless, overlapping rCpGms from our RRBS-based TCX also had high correlations with those from the other array-based datasets (Spearman Coefficients: 0.74 – 0.828) (**Figure S6**). Thus, while not identical to our DNAm measurements, rCpGms from these other AD brain cohorts could serve as replication datasets for our findings.

For replications, we focused on the 738 TCX rCpGms that are significantly associated with brain levels of both Total tau_TBS_ and pTau_TX_ (**Figure 3**). These 738 rCpGms are high confidence both due to their FDR-significant and biologically congruent directions of associations with two brain biochemical AD endophenotypes. Of these 738 rCpGms, 600 were present in at least one of the replication datasets. Although BDR and ROSMAP did not have brain biochemical AD-related protein measurements, they both have Braak stages, which is a measure of tau pathology. To consider an rCpGm as having a biologically congruent association in the replication datasets, the direction of effect would be the same for Braak stage and pTau_TX_ and opposite for Total tau_TBS_. This is based on prior work^9^ and positive correlations of Braak stage with pTau_TX_ (Spearman correlation ρ = 0.289, pval = 3.72E-10), but negative with Total tau_TBS_ (Spearman correlation ρ= -0.30, pval = 6.29E-11) in our dataset (**Figure S7**).

Using these criteria, 93 out of 600 tested rCpGms had nominally significant (p-value ≤ 0.05) and biologically congruent associations with Braak stage in BDR and/or ROSMAP (**Figure 4**, **Table S8, Table S9**). Of these 93, 3 replicated in all 3 datasets (tier 1), 23 in 2 of the datasets (tier 2), and 67 in 1 of the replication datasets (tier 3). We further tested the association of these rCpGms with Thal phase and AD diagnosis in the replication datasets and also found biological congruence of associations across these additional AD measures (**Figure 4**, **Table S8, Table S9**). In other words, rCpGms with FDR-significant negative associations with Total tau_TBS_ and positive associations with pTau_TX_ in our dataset, had positive associations with Braak, Thal and AD diagnosis in one or more of the replication datasets and vice versa.

**Figure 4:**
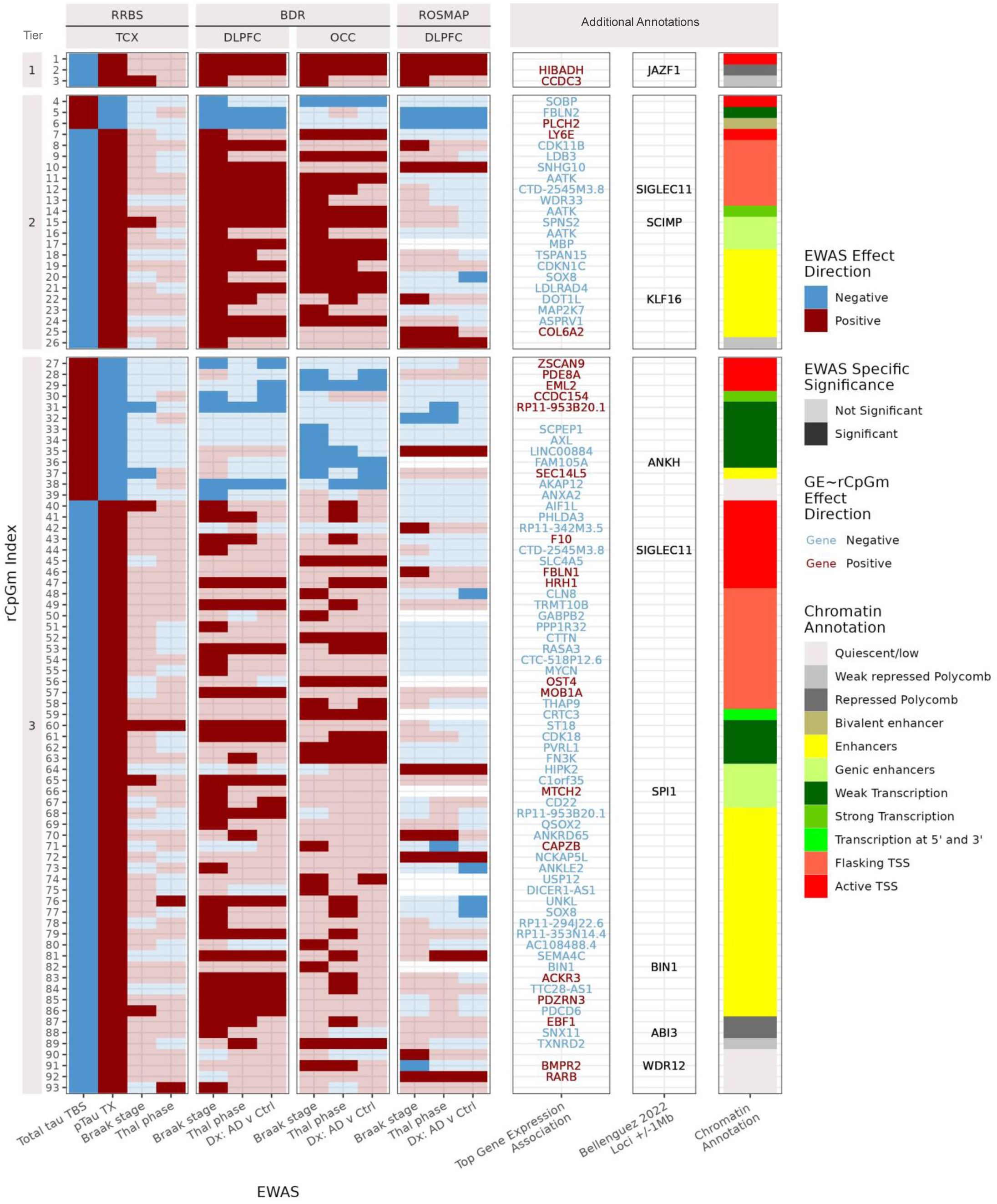
Replication of FDR significant brain tau-associated rCpGms in independent datasets. The y-axis depicts the 93 rCpGms that are FDR significant (FDR pval < 0.05) in the Mayo Clinic RRBS-TCX total tau_TBS_ and pTau_TX_ EWAS and have opposite directions of associations with these tau endophenotypes. These rCpGms are nominally significant (p-value < 0.05) and have congruent direction of associations with Braak stage in ROSMAP and/or BDR. Tier 1 rCpGms (N= 3) are associated with Braak stage in all three replication datasets (BDR-DLPFC, BDR-OCC, and ROSMAP-DLPFC); Tier 2 (N=23) are associated with Braak stage in two of the replication datasets; Tier 3 (N=67) are associated with Braak stage in one of the replication datasets. Also included are additional associations of the rCpGms with Thal phase and AD diagnosis (Dx) with significant associations defined as those with a p-value < 0.05. Non-colored associations in ROSMAP indicate that this region was not measured on the array. Each rCpGm is additionally annotated with the top FDR significant gene expression association (GE∼rCpGm) colored by direction of effect, their nearest known AD risk loci (+/- 1Mb, Bellenguez 2022), and by their chromatin state annotation from the Roadmap Epigenomics 15 chromatin state model. See supplemental data **Table S12** to connect rCpGm Index numbers with rCpGm positional data. Red indicates a positive direction of effect, blue indicates a negative direction of effect, and the transparency indicates if the association passed nominal significance threshold. **Dx** = Diagnosis; **FDR** = false discovery rate; **N** = Number; **pTau** = phospho-tau; **TBS** = soluble tris buffered saline fraction; **TX**= membrane associated Triton-X fraction

When we annotated these 93 rCpGms for proximal (+/- 1Mb) variants identified in AD genome-wide studies^53^, we found that a number of them reside near known AD risk loci including *JAZF1, SIGLEC11, SCIMP, KLF16, BIN1, SPI1, ABI3, ANKH, WDR12, and SIGLEC11*^53^ (**Figure 4**). Collectively, these results support independent replication of our 93 high confidence TCX rCpGms for their association with brain tau-related and other AD endophenotypes, which may also have implications for AD risk.

### Tau related rCpGms associate with expression of nearby oligodendrocyte and myelin related genes

Most of the FDR significant AD endophenotype associated rCpGms in our dataset are located in active chromatin regions (**Figures 3-4**). To further characterize the associations between these rCpGms and brain gene expression levels, we leveraged bulk CER and TCX RNAseq data from the same individuals. After RNAseq QC, we paired rCpGms to genes +/- 500kb from the start and end of each rCpGm yielding 1,612,499 TCX and 1,834,887 CER *cis*-rCpGm-gene pairs. We then tested the association between *cis*-rCpGm-gene expression pairs using linear regression.

We identified 12,973 TCX and 1,413 CER FDR significant *cis-*rCpGm-gene pairs (FDR p-value ≤ 0.05) (**Table S10, Table S11**). Many of the significant *cis*-rCpGm-gene pair associations had rCpGms annotated to expression regulation terms including active TSS, flanking active TSS, and enhancer sites (51% in TCX and 32% in CER). Genes previously reported to have AD-related methylation changes nearby, including *RHBDF2*^30,31^, *IRF8*^31^, *CCND1*^31^, and *SPI1*^32^ had both FDR significant rCpGm associations with AD endophenotypes and *cis*-rCpGm-gene expression associations in our dataset (**Table S10, Table S11**).

To further annotate the 93 high-confidence tau and other AD-endophenotype associated rCpGms, we tested their corresponding 1,464 *cis-*rCpGm-gene expression pairs (1,225 unique genes). Many of these 93 rCpGms reside in more active chromatin regions such as Active TSS, Flasking TSS, or Enhancers, suggesting their functional consequences in gene expression (**Figure 4**). Indeed, when tested for their effects on brain gene expression, these rCpGms had FDR significant associations with nearby genes measured from the same donors (**Tables S10-S12**). There were 535 significant *cis-*rCpGm-gene expression associations (459 unique genes) which included previously implicated AD genes like *BIN1*^53–55^*, ANXA2*^56,57^, CDK18^56,58,59^, *AKAP13*^60^*, RORA*^61,62^*, KIFC1*^63^*, and KLK6*^64,65^(**Table S12)**.

To determine if this set of 459 genes, which correspond to the 93 tau-related high confidence rCpGms, implicates common pathways or cell types, we performed gene set enrichment analysis. The top three nominally enriched gene ontology terms for these genes relate to positive regulation of myelination (GO:0031643, pval = 3.4E-5), gliogenesis (GO:0042063, pval = 1.4E-4), and microtubule nucleation (GO:0007020, pval = 1.5E-4). Consistent with these GO terms, this gene set was also enriched for oligodendrocyte marker genes. Using the top 1000 BRETIGEA^66^ cell type markers genes, there were 45 oligodendrocyte genes amongst these 459 (p-value = 9.8E-05), 11 of which were also in the top 100 BRETIGEA^66^ markers (p-value = 4.0E-05). These top 11 oligodendrocyte marker genes with rCpGm associations are *MAG, KLK6, ST18, MYRF, LDB3, MBP, CD22, TGFA, RTKN, SOX8,* and *CLMN.* We next investigated these 11 genes and an additional gene *CKD18* further in both our and external datasets. Although part of the 1000, and not the 100 top oligodendrocyte marker genes^66^, *CKD18* was previously implicated with tau^59^ and AD DNAm changes^67^, thereby leading us to include this gene in additional analyses.

All 12 oligodendrocyte genes have negative associations of their brain expression and paired rCpGm methylation levels (**Figure 5**). There were 16 rCpGm-gene expression pairs for these 12 oligodendrocyte genes and 15 unique rCpGms. All 15 rCpGms had FDR significant negative associations with Total tau_TBS_ and positive associations with pTau_TX_ (**Figure 5**). These rCpGm∼tau endophenotype associations remained significant even after adjusting for oligodendrocyte cell proportion estimates (**Table S13**), indicating that they are not driven by changes in proportions of oligodendrocytes in bulk brain tissue. These rCpGms had additional nominally significant associations with other AD-related endophenotypes, the most consistent of which were negative associations with APOE_TX_ levels and positive associations with pTau_FA_ (**Figure S8**). Almost all of the individual CpGms that comprise these oligodendrocyte rCpGms have the same direction of association with Total tau_TBS_ or pTau_TX_ as their corresponding rCpGm, although few individually reach FDR significance (**Figure S9**). This again underscores the superior power of our novel and biologically relevant rCpGm in comparison to individual CpGm analysis in detecting DNAm associations.

**Figure 5:**
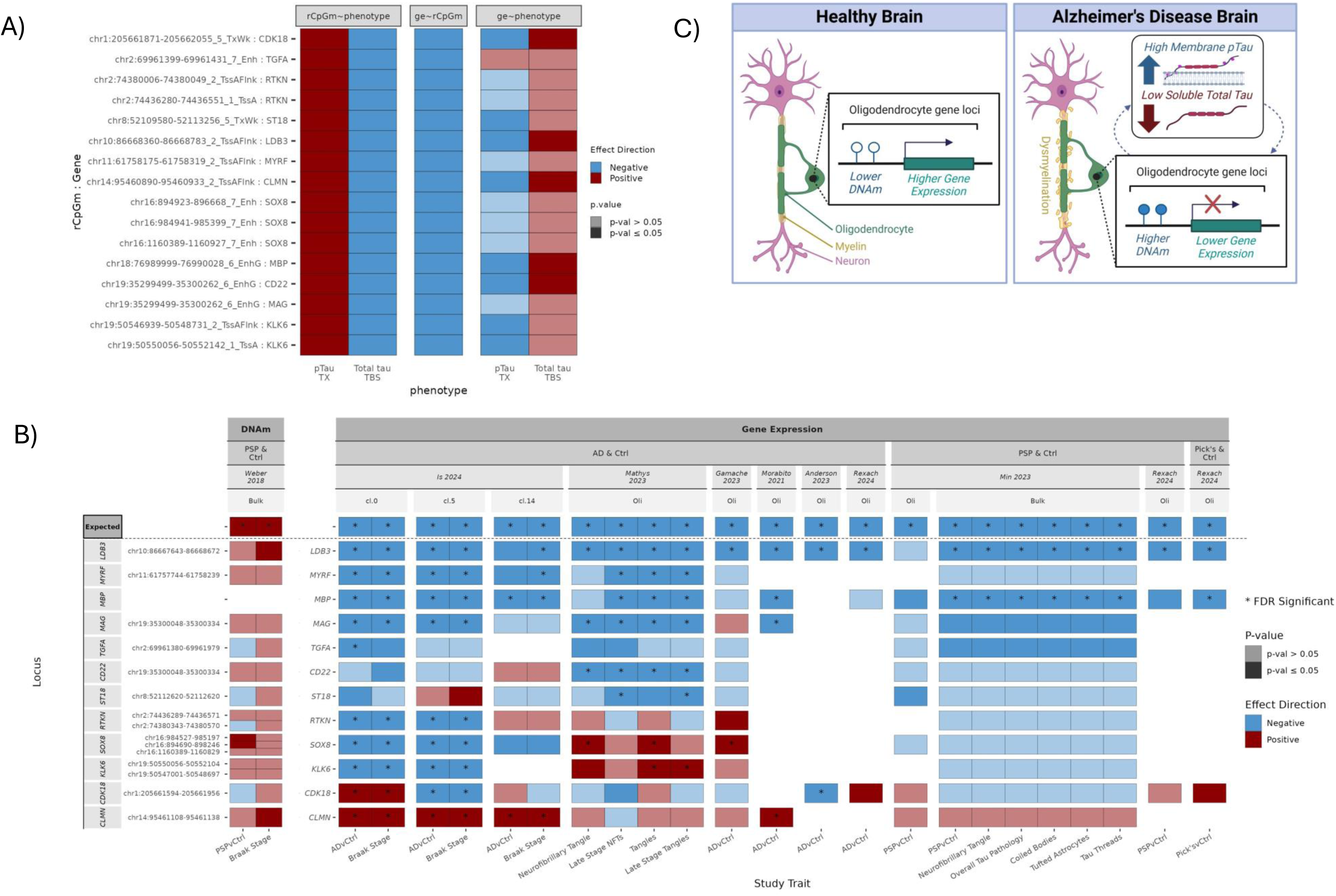
Top rCpGm and oligodendrocyte gene loci consistently associate with tau-related phenotypes across studies. A) Association results of the top oligodendrocyte specific rCpGm-gene pairs related to TCX tau levels in our dataset across three models: 1) rCpGm ∼ tau phenotype, 2) Gene expression ∼ rCpGm levels, 3) Gene expression ∼ tau phenotype. Red indicates a positive association and blue indicates a negative association. B) Association results of top tau-related rCpGm or oligodendrocyte gene expression levels to multiple tau related endophenotypes pulled from multiple, recently published studies. These studies include Weber 2018 (N= 90 PSP, N = 66 Control), Is 2024 (N = 12 AD, N = 12 Control), Mathys 2023 (N = 238 AD, N = 189 non-AD), Gamache 2023 (N = 12 AD, N = 12 control), Morabito 2021 (N = 11 AD, N = 7 control), Anderson 2023 (N = 7 AD, N = 8 Control), Rexach 2024 (N = 10 AD, N = 11 PSP, N = 10 Pick’s, N = 10 Control), and Min 2023 (snRNAseq: N = 18 PSP, N = 16 Control; Bulk: N = 281 PSP, N = 127 Control). Top X-axis is faceted in the first levels between whether DNAm or gene expression levels were tested, second the disease type of the study cohort examined, third the study examined, fourth whether bulk DNAm/RNAseq data, pseudobulk oligodendrocyte clusters from sn/scRNAseq data or specific oligodendrocyte assigned clusters were analyzed, and finally the bottom x-axis is the study trait investigated. The first row “Expected” depicts whether we expect a positive (red) or negative (blue) association based on results from our dataset. Locus annotations on the y-axis include the rCpGms defined by CpGms present in the Weber *et al* 2018 study that overlap with our TCX rCpGms of interest in panel A. White areas indicate a result for that association was not reported from the study indicated. C) Hypothesized relationship between TCX tau levels, DNAm, and oligodendrocyte gene expression in the AD brain. We hypothesize that in the healthy brain, there are lower levels of DNAm at these loci allowing oligodendrocytes to express implicated myelin related genes, but in the AD brain, as pTau_TX_ levels increase and total tau_TBS_ levels decrease, higher levels of DNAm are introduced to these regions which in turn decreases expression of the oligodendrocyte and myelin related genes potentially leading to oligodendrocyte dysfunction and dysmyelination. Created in BioRender.com **abs** = absolute value; **AD**= Alzheimer’s Disease; **cl.** = cluster; **Ctrl** = Control; **DNAm** = DNA methylation; **ge** = Gene expression; **Oli** = Pseudobulk Oligodendrocytes; **PSP**= Progressive Supranuclear Palsy; **pTau** = phospho-tau; **p-val** = p-value; **TBS** = soluble tris buffered saline fraction; **tTau** = total tau; **TX**= membrane associated Triton-X fraction

We further evaluated the top oligodendrocyte genes with tau-related rCpGm associations using our dataset and independent studies. Within the same brain donors, TCX expression levels of these oligodendrocyte genes also had inverse directions of associations with TCX levels of Total tau_TBS_ and pTau_TX_. Of these 12 top oligodendrocyte genes, 5 nominally associated with both Total tau_TBS_ and pTau_TX_ levels (*CDK18, LDB3, MBP, CD22, CLMN*) and 2 with pTau_TX_ (*KLK6, ST18*) in the TCX **(Figure 5A, Table S14**). All but one of these associations (*TGFA*-pTau_TX_) had directions of effect that are biologically congruent with both rCpGm-tau and rCpGm-gene associations. Notably, *TGFA*, unlike the other marker genes, is not only expressed in oligodendrocyte cells but is also expressed in oligodendrocyte precursor cells which may underlie this discrepancy. Upon adjustment for oligodendrocyte cell proportion estimates, *CDK18, MBP,* and *CD22* remained nominally associated with both Total tau_TBS_ and pTau_TX_ levels with consistent effect directions (**Table S14**).

Using three independent brain bulk RNAseq datasets of AD and control brains, we next investigated the association of these top oligodendrocyte genes with Braak stage in the Mayo TCX^68^, Mount Sinai BM22^69^, and ROSMAP^70^ datasets from the AMP-AD study^71^. Expression of *LDB3* was associated with Braak stage in the Mayo TCX (beta = -0.19; p-value = 8.47E-3) and ROSMAP (beta = -0.058; p-value = 7.22E-3) datasets with a biologically congruent negative direction of effect (**Table S15**). Using AMP-AD data, *MYRF* and *CDK18* have also been nominated as potential therapeutic targets for AD^72^, providing additional support for a role of these oligodendrocyte genes in tau-related outcomes.

To determine whether these brain rCpGm-oligodendrocyte gene-tau associations are unique to AD vs. conserved across other tauopathies, we investigated available data from methylation or gene expression studies of progressive supranuclear palsy (PSP) and Pick’s disease. Further, we analyzed oligodendrocyte cell clusters from published brain single nucleus datasets of tauopathies to establish these associations in the cell type of interest. There was one brain DNAm study conducted from the prefrontal cortex of PSP and control donors^73^ to which we applied our QC, rCpGm method and conducted an EWAS of PSP risk and Braak stage (90 PSP and 66 controls) (**Tables S3, S16, Figure 5B**). We determined biologically congruent associations with Braak stage across the 12 top oligodendrocyte rCpGms that also had consistent PSP risk associations for all but three of the genes (*TGFA, ST18, CDK18*). This finding supports a model where rCpGm-oligodendrocyte gene-tau associations are not unique to AD but may be a shared disease mechanism across tauopathies.

We next analyzed all available single nucleus RNAseq (snRNAseq) data from sizeable brain datasets of AD and other tauopathies^74–80^ to replicate the top oligodendrocyte gene expression associations with disease risk and tau endophenotypes in an oligodendrocyte cell-type specific manner. These independent snRNAseq datasets and brain regions were from Is et al.^74^ (24 TCX, AD and controls), Mathys et al. (427 prefrontal cortex, AD and controls)^75^, Gamache et al. (24 TCX, AD and controls)^76^, Morabito et al. (18 prefrontal cortex, AD and controls)^77^, Anderson et al. (15 dorsolateral prefrontal cortex, AD and controls)^78^, Rexach et al. (41 insular cortex, AD, PSP, Pick’s disease and controls)^79^, and Min et al. (34 TCX, PSP and controls)^80^. In addition to these snRNAseq data from a total of 583 independent brain samples from multiple regions and diagnoses, Min et al.^80^ also had bulk RNAseq data from 408 PSP and control TCX samples. We tested the top brain rCpGm associated oligodendrocyte genes for their oligodendrocyte-specific expression associations with risk for AD^74–79^, PSP^79,80^, Pick’s disease^79^, tau-related (**Figure 5B**) and other available endophenotypes (**Figure S10**) in these datasets. Despite differences in the cohorts, brain regions and diagnoses, there was remarkably consistent association for oligodendrocyte-specific expression for many of these top 12 genes with these outcomes in a direction that would be expected based on the rCpGm-oligodendrocyte gene-tau associations we identified. Most of these genes had lower expression levels in AD and other tauopathies and were negatively associated with tau-endophenotypes such as Braak stage and neurofibrillary tangles. Most consistent associations were observed for *LDB3, MYRF, MBP, MAG, TGFA,* and *CD22* loci where the implicated rCpGms positively associate with pTau_TX_, negatively with Total tau_TBS_ and brain gene expression in our data (**Figure 5A**, **Tables S12-14**). These rCpGms associate positively with PSP risk and Braak in Weber et al.^73^ DNAm data (**Figure 5B**, **Tables S16)**. Further, oligodendrocyte cluster specific expression of these genes associate negatively with tauopathy risk and tau traits^74–80^ (**Figure 5B**, **Figure S10**).

To determine whether genetic variants could account for the tau-related DNAm changes observed for these top oligodendrocyte gene related rCpGms, we performed a *cis*-methylation quantitative trait locus (mQTL) analysis between these rCpGms and imputed genetic variant dosages within a distance of +/- 500kb. There were 160 FDR significant SNP-rCpGm associations (**Table S17**). The rCpGms implicated by the mQTL associated with *ST18* (14 SNP-rCpGm pairs), *MBP* (122), *CLMN* (2), *MAG* & *CD22* (2), *KLK6* (1), and *RTKN* (19) gene expression. Of these 160 unique SNPs, 13 near *MBP* nominally (p-value ≤ 0.05) associated with Total tau_TBS_ levels, the top 2 of which were also trending towards association (p-value < 0.052) with pTau_TX_ levels (rs13381407, rs12457073: R^2^ = 1, D’=1)^9^ (**Table S17)**, but none associated with AD risk^53^. These results suggest that genetic variation is unlikely to account for the rCpGm-gene-tau relationships at these loci, with the possible exception of the MBP locus.

Collectively, these findings support a model for tauopathies such that with increasing brain levels of tau neuropathology, pTau_TX_, and decreasing Total tau_TBS_, DNA hyper-methylation occurs in oligodendrocyte/myelin gene regions leading to reduced expression of these genes and potentially downstream oligodendrocyte dysfunction and dysmyelination (**Figure 5C**).

## Discussion

AD is a complex and heterogeneous disease with variability not only in its clinical presentation but also neuropathological features and brain biochemical levels of AD-related proteins^5–8^ like Aβ, tau, and apolipoprotein E (APOE). Capturing the genomic basis of this variability can reveal novel molecular mechanisms of AD and identify causal factors that underlie disease heterogeneity. These are essential to devise precision therapies for this neurodegenerative disease much like is being done in other complex and heterogeneous diseases, like cancer^81^. We^82^ and others^83^ have utilized deep AD neuropathology endophenotypes in GWAS to uncover genetic variants associated with variability of these neuropathologic measures. Previous brain methylation studies in AD focused on associations with clinical features of this disease^30–32,38,39^, with fewer studies focusing on AD neuropathology^32,84^. We previously identified genetic variants that may influence variability in brain biochemical measures of AD-related proteins^9^, demonstrating their significant potential as deep endophenotypes in AD genomic studies. However, to our knowledge, there are no studies that previously explored the combined genetic, epigenetic and transcriptomic landscape of both AD neuropathology and brain biochemical measures.

This study employed a deep endophenotyping approach to identify important molecular mechanisms of DNAm that influence levels of neuropathology and biochemical states of core AD related proteins in Alzheimer’s Disease brain tissue. By leveraging the Roadmap Epigenomics 15-chromatin state model to group individual CpGm sites into rCpGms, we established a novel methodology to investigate grouped CpGms in a biologically relevant manner increasing our ability to identify robust associations with these endophenotypes. Through our innovative methodology to investigate regional CpGm, EWAS, and integrative analyses, we found that although all the tested endophenotypes are core to AD pathophysiology, each have distinct CpGm architectures underlying their variability in the AD brain. This establishes the role of epigenetic mechanisms in influencing heterogeneity of these AD brain endophenotypes (**Figure 1**).

We found highly consistent and biologically congruent patterns of average rCpGm levels across three independent datasets from Mayo Clinic (this study), ROSMAP^30^ and BDR^32^, collectively comprising 2,515 samples representing four different brain regions (TCX, CER, DLPFC, OCC) (**Figure 2**, **Supplementary Figure S3**). The striking similarities in the rCpGm levels and patterns, despite variabilities in cohort, brain region, DNAm measurement technologies and methylation coverage in these cohorts underscores the strong biological underpinning of our novel rCpGm method and supports its broad utility in analyzing regional levels of DNA methylation across measurement platforms, tissue types, and disease states.

Using our rCpGm grouping approach, we identified 5,478 FDR significant TCX rCpGms where the majority (99.7%) associated with brain tau levels, i.e. total tau_TBS_ and/or pTau_TX_ (**Figure 3**). We found 738 rCpGm associations with shared but opposite directions of effect between Total tau_TBS_ and pTau_TX_ suggesting that there are shared DNAm related mechanisms that inversely impact levels of soluble total tau and membrane-bound phosphorylated tau (pTau) in the AD brain. These findings may have broader implications for therapeutic development as targeting these shared pathways may influence levels of both brain Total tau_TBS_ and pTau_TX_. Conversely, similar to previous AD CpGm studies, we found no FDR significant associations of CER rCpGm with neuropathology^31,39,85^, although there were some individual CER CpGm associations, but to a much lesser extent than those for TCX CpGm. This may suggest that CER, which has less AD neuropathology, may consequently have less DNAm perturbations. Together, these results demonstrate the effectiveness of using a deep endophenotyping approach combined with integrative multi-omics to identify important and phenotype specific molecular mechanisms that likely contribute to the pathophysiology of AD.

The finding that the largest number of significant brain rCpGm associations occur with brain tau related measures suggests that epigenetic dysregulation may occur more within the context of tau rather than amyloid in the AD brain. This parallels a previous epigenetic study which suggested that large-scale changes in chromatin states marked by H3K9ac, a typical open chromatin marker, occur as a result of tau rather than Aβ burden^28^. Interestingly, these tau related epigenetic associations juxtapose our previous GWAS findings in this same dataset where we identified that genetic variants associate with brain amyloid rather than tau proteins^9^. In sum, these findings may suggest a model where distinct AD-related proteins in the brain may be associated with different genomic perturbations. Specifically, genetic variants primarily drive brain biochemical measures of Aβ, whereas brain methylation changes associate with brain tau levels. These results demonstrate how integrating multi-omics measures with distinct deep AD phenotypes can begin to unravel the molecular basis of different facets of AD, although the precise genomic mechanisms underlying this distinction remain to be uncovered.

The regional DNAm associations we found with brain tau endophenotypes in our Mayo Clinic cohort were replicated in the two large, independent datasets of AD and control brains from the ROSMAP^30^ and BDR^32^ cohorts (**Figure 4**). As these replication cohorts do not have brain tau biochemical measurements, we used Braak stage as a tau neuropathology endophenotype. Remarkably, we found that of the 600 tau-related rCpGms in our cohort, which were also available from these replication datasets, 93 showed significant and biologically congruent associations in BDR and/or ROSMAP. Ten of these replicated rCpGms are located near (+/- 1MB) known AD risk GWAS loci^53^ suggesting an epigenetic mechanism in conferring AD risk at these loci. Characterization of the 93 replicated rCpGms by integrating matched brain gene expression measures from the same brain donors revealed a likely functional role of these rCpGms on expression of nearby genes. Many of these genes also have previous connections with tau, such as *BIN1* which is known to interact with tau proteins in extracellular vesicles^86,87^ including tau Thr231^88^ which is the pTau species measured in our study, *ANXA2*^56,57^, as well as *CDK18*^58,59^ and *AKAP13*^60^ which can phosphorylate tau.

Further interrogation of the genes associated with the tau-related rCpGms revealed enrichment in oligodendrocyte marker and myelination-related genes and GO terms. Importantly, higher brain levels of pTau_TX_, which is considered to be a toxic tau species^9,89^ is associated with hypermethylation at the rCpGms near the oligodendrocyte genes and biologically consistent lower brain gene expression levels for many of them. Downregulation of oligodendrocyte gene expression in AD brains and other tauopathies have previously been reported by our group^80,90,91^ and others^92^. Our integrated brain methylation and gene expression results reported here suggest that the transcriptional downregulation of oligodendrocyte and myelin related genes in tauopathies may be mediated by pTau related CpG hypermethylation near these genes.

We further characterized and validated the rCpGm and gene expression associations for these oligodendrocyte genes using additional, independent data generated by both our group^74,80^ and others^73,75–79^. Considering our Mayo Clinic discovery, ROSMAP^30^ and BDR^32^ replication cohorts in this study together with these additional validation cohorts, these top oligodendrocyte genes were assessed in 2,671 rCpGm, 1,080 bulk RNA and 583 snRNAseq brain sample measurements representing 8 different brain regions (**Figure 5**). Of the oligodendrocyte genes thus characterized, *LDB3, MBP*, *MAG*, and *MYRF* had the most consistent brain rCpGm, bulk or single nucleus gene expression associations with disease risk or tau endophenotypes across AD and other tauopathy cohorts.

*LDB3* encodes LIM Domain Binding 3 (LDB3) which is a protein that interacts with the cytoskeleton and plays a role in targeting and clustering membrane proteins often studied in muscle tissues, but it is also expressed in the central nervous system^93,94^ and oligodendrocytes in the brain^95^ (proteinatlas.org/ENSG00000122367-LDB3)^96^. Although genetic variants in *LDB3* are often implicated in myopathies^93,97,98^, there are suggestive associations with AD risk in Caribbean Hispanics^99,100^ and differential expression in AD^76,101^. There is also evidence that *Ldb3* decreases in a rat model of demyelinating epilepsy^102^ and expression of *LDB3* is downregulated in Huntington’s disease human brain tissue and mouse models^103^. In our study, brain *LDB3* gene expression and its rCpGm are associated with brain tau endophenotypes, AD and other tauopathies. Like *LDB3*, tau is a cytoskeletal protein that interacts with and likely stabilizes microtubules^89,104^. This raises the possibility that these two proteins may interact through cytoskeletal activity and collectively contribute to maintenance of oligodendrocyte and neuronal activity in health or their demise in the presence of toxic tau species (**Figure 5C**). In addition, *MBP, MAG,* and *MYRF* encode important factors for myelination. Myelin basic protein (MBP) and myelin associated glycoprotein (MAG) are major components of myelin^105^, and myelin regulatory factor (MYRF) is a transcription factor regulating oligodendrocyte differentiation and myelin-related gene expression^106^. Previous cell culture models showed that knockdown of tau mRNA led to a decrease in protein levels of MBP, disruption of microtubule formation, and disturbance of myelin sheath formation^107^. This finding is aligned with our results suggesting that total tau may have a protective role in maintaining oligodendrocyte epigenetic and transcriptional homeostasis, whereas pTau has the opposite, toxic effect (**Figures 3-5**).

A relationship between myelination and tau is supported by many previous studies by our group and others^107–113^ including downregulation of myelination related gene expression networks in AD as well as the primary tauopathy of PSP. There are white matter abnormalities and myelin loss in human AD patients^114–116^ and mouse models of tauopathy^117–119^. Further, higher levels of tau-PET correlate with lower myelin levels^120^. Importantly, oligodendrocytes can propagate tau pathology in the absence of neuronal transmission, suggesting a potential role of the myelin sheath in pathologic tau propagation^121^. Primary tauopathies, like PSP, can present with pathological tau inclusions in oligodendrocytes^122^. Previous studies of mice expressing human mutant tau (P301L) exclusively in oligodendrocytes found evidence that oligodendrocytic tau inclusions preceded myelin and neuronal axon disruption and that they may in part cause neurodegeneration^123^.

Our epigenome-wide association study of deep brain AD protein biochemical levels, coupled with our novel rCpGm approach, and integrated transcriptome and epigenome data from >4,000 collective brain samples demonstrate strong, replicable and biologically consistent associations between brain tau and hypermethylation near oligodendrocyte genes likely leading to their lower expression, dysmyelination and neurodegeneration in AD and other tauopathies (**Figure 5C**). While future studies are needed to precisely delineate the functional mechanisms behind these results, previous studies also provide evidence for these conclusions. Expression levels of *BIN1*, an AD risk gene^23,53,55^, associate with AD and there is loss of *BIN1* expression and demyelination in multiple sclerosis lesions^124^. *BIN1* also emerged as a gene with brain tau, rCpGm and expression associations in this study. Others have shown hypermethylation of individual CpGms near *MBP* in AD brain^37,84,125,126^ and plasma^127^ as well as in multiple sclerosis lesions^128^, and suggested that Braak associated CpGms in bulk brain tissue likely reflect changes in glia cells including oligodendrocyte enriched populations^32^. Moreover, the importance of DNAm in oligodendrocyte differentiation and function has been appreciated where disruption of normal DNAm mechanisms can cause oligodendrocyte and myelin dysfunction^129,130^. These observations support the connection between DNAm and tau in oligodendrocyte function and may open up potentially important opportunities for therapeutic interventions for neurodegenerative diseases^131^ targeting correction of disrupted myelination as well as DNAm mechanisms.

Our study has multiple strengths including our discovery^9^ and replication^30,32^ cohorts of AD and control donors with DNAm measures from four brain regions of 2,515 samples which we analyzed using our novel rCpGm analytic approach. Mayo Clinic discovery cohort has deep endophenotypes of AD neuropathology and precise biochemical measures of core AD proteins. In this cohort, we generated DNAm measures via RRBS, which allowed us to investigate close to two million high confidence CpGm sites, more than doubling the number of sites that can be investigated by the frequently used array platforms. We developed a novel, biologically relevant methodology to investigate DNAm on a regional, rather than individual CpGm level. We determined that our regional grouping method can be applied to DNAm data generated with different technologies, yielding remarkably consistent rCpGm levels across independent datasets and tissue types demonstrating broad utility of our innovative analytic approach. Additionally, investigating DNAm grouped by chromatin regions^48^, rather than arbitrary groupings or individual CpGm, has better functional alignment providing biologically relevant results. We integrated additional omics measures from the same donors including brain gene expression and genotype data and identified 93 rCpGms corresponding to 535 genes that have replicable and biologically congruent associations with brain tau levels. These associations are enriched for myelination pathways and genes, which we further validated using additional brain methylation, bulk and single nucleus expression data from AD and other tauopathy cohorts totaling >4,000 samples. Our results uncovered opposite roles for brain total tau (likely beneficial) and ptau (detrimental) and support a model where brain tauopathy influences rCpGm resulting in hypermethylation and reduced expression of oligodendrocyte genes culminating in dysmyelination.

Despite these strengths, there are some limitations to our study. As with many DNAm measurement approaches, RRBS cannot differentiate between 5mC and 5hmC methylation, with the latter found at higher levels in central nervous system tissue types compared to other tissue types^132,133^. While this should not affect our highly replicable findings, we may have missed some specific methylation associations of either 5mC or 5hmC, as they can have opposite functional consequences^134–138^ including in AD^29,139–142^. Additionally, we extracted CER DNA using a manual rather than the automated approach used for TCX DNA, to achieve similar quality DNA from both tissue regions. Although our pilot studies (see **Methods**) demonstrate no significant contributions to CpGm variation by DNA extraction method, we cannot entirely rule out this potential for all CpGm sites. Our study is conducted on bulk brain tissue, and individual cell types can have different CpGm compositions^32,143^. Although the associations for the top hits we identified remained even after adjusting for neuronal proportions, future single cell type specific DNAm studies, particularly those from glial cell types^32^, will be important in parsing out potential cell type specific rCpGm effects in these associations. Lastly, given the lack of brain multi-omics data from diverse populations, the cohorts in this study are from non-Hispanic white brain donors. This is a major gap in the field, which we anticipate will be addressed with foundational brain multi-omics data recently emerging from multi-ethnic brain donors^144^.

In conclusion, the landscape of regional DNAm integrated with gene expression reveal a distinct epigenomic architecture underlying variability of brain tau biochemical species and neuropathology in AD. Our robust and replicable findings suggest that tau-related epigenetic dysregulation of gene expression impacts oligodendrocytes and may be a critical mechanism of dysmyelination in AD and other tauopathies. Our study identifies novel genes implicated in AD, such as *LDB3*, and many others that are replicated across multiple independent datasets. This study demonstrates the potential of applying rigorous and innovative multi-omics analysis to deep endophenotypes to unravel the complex heterogeneity of AD, which also has implications for other neurodegenerative diseases.

## Methods

### Brain Samples

Post-mortem superior temporal gyrus cortex (TCX) and cerebellum (CER) tissue samples were collected by the Department of Neuroscience, Mayo Clinic Brain Bank (Jacksonville, Florida, USA). These samples are a part of the Mayo Clinic AD-CAA (MC-CAA) study, previously described^9,82^. All samples had a confirmed AD neuropathological diagnosis with a Braak stage ≥ four and Thal phase ≥ two, were scored for cerebral amyloid angiopathy (CAA) pathology, had an age at death greater than 55 years, and due to availability, were all non-Hispanic white (**Table S1**). Donors with co-pathologies were not excluded to allow for inclusion of the full spectrum of neuropathologies in AD. Braak stage, Thal phase, and CAA scores were measured using previously established protocols^3,4,10,145,146^. Intermediate Braak stages were grouped with the next lowest stage as follows, stage 3.5 is 3, 4.5 is 4, and 5.5 is 5 as detailed previously^10,82^ (**Table S1**, **Figure S1**). This study was approved by the appropriate Mayo Clinic Institutional Review Board.

### Biochemical Measures

Biochemical measures from a subset of superior temporal cortex brain samples included the MC-CAA study described previously^9,10^ were utilized for this study. Biochemical measures include five AD-related proteins (APOE, Aβ40, Aβ42, tau, and phospho-tau (Thr231)) from three tissue fractions (soluble, membrane-bound, insoluble). Briefly, supernatant fractions were collected after three sequential buffer treatments of tissue homogenate and resulting pellets: first with tris-buffered saline buffer (TBS), second with detergent (TBS/1% Triton X) buffer (TX), and lastly with formic acid buffer (FA). Protein quantification was performed via ELISA for each fraction and normalized against total protein quantities. All biochemical measures were transformed by either the natural log or square root to approximate a normal distribution including the Aβ40/42 ratio. Transformed measures were then standardized with a mean of zero and standard deviation of one using the z-score formula to allow for comparisons of effect sizes between biochemical measure analyses. Of the 469 MC-CAA samples with biochemical measures, 452 also had DNA methylation data (**Figure S1**).

### Genotype Data

Genome wide genotypes (GWG) were generated in previous studies and are described in detail elsewhere^9,82^. Briefly, DNA was isolated for GWG from brain tissue using the AutoGen245T instrument according to manufacturer’s instructions including an RNase A (Qiagen) digestion. GWG were collected using the Infinum Omni2.5 Exome 8 v1.3 genotyping array, formatted in PLINK (v1.9)^147,148^ and rigorously quality controlled (QC). Variants passing QC were imputed to the haplotype reference consortium (HRC) keeping only variants with an imputation quality R2 ≥ 0.7 and minor allele frequency (MAF) ≥ 2%. Variant base pair (BP) positions were lifted from hg19 to hg38 using the *liftOver* package and corresponding chain files provided by UCSC Genome Browser. After mapping coordinates in hg19 to hg38, positional information in the plink files were updated using the ‘--update-map’ flag. Of the N = 6,724,981 imputed and quality controlled genetic variants aligned to the hg19 reference genome, N = 6,721,677 successfully lifted over to the hg38 reference and could be analyzed.

### DNA methylation

DNA from 40-70 mg of TCX tissue was isolated using the AutoGen245T instrument according to manufacturer’s instructions including an RNase A (Qiagen) digestion prior to loading. The CER samples failed DNA isolation using the AutoGen instrument, so DNA was isolated from 10-15 mg of CER tissue using the same AutoGen Reagent kit and instrument steps manually. Pilot experiments were performed comparing TCX DNA isolated in triplicate from 3 samples using the AutoGen instrument vs manual methods as well as 2 CER samples in triplicate manually to determine if extraction method significantly impacted overall DNA methylation (DNAm) levels (**Figure S11**). Following DNA extraction of all samples, Reduced Representation Bisulfite Sequencing (RRBS) was performed on 25 ng of genomic DNA for each sample by the Mayo Clinic Genome Analysis Core (GAC) in Rochester, MN following manufacturer’s recommendations for the NuGen Ovation RRBS Methyl-Seq System 1-16 (Tecan Genomics). The EZ DNA Methylation Kit (Zymo Research) was used for bisulfite conversion. RRBS libraries were sequenced on the HiSeq 4000 (Illumina) with 8 samples per lane and 50 bp paired end reads. The Streamlined Analysis and Annotation Pipeline for Reduced Representational Bisulfite Sequencing (SAAP-RRBS)^149^ was used to align reads to the hg38 reference genome and to get CpGm ratio information. Briefly, FASTQs were trimmed to remove diversity and adaptor sequences using cutadapt, and any reads with less than 15bp were discarded. Trimmed FASTQs were then aligned against the hg38 reference genome using BSMAP^150^. Samtools^151^ was used to get mpileup and custom PERL scripts were used to determine CpG and non-CpG methylation levels along with bisulfite conversion ratios. All samples had a conversion ratio >98% and we kept only CpG sites with at least 10x coverage. One triplicate TCX sample (*TCX_Autogen_1c*) had low read coverage and so was not included in the downstream comparisons (**Figure S11a**). In a Principal Component Analysis (PCA) of the top 50% most variable CpG sites with 10x coverage (N= 535,165 CpG’s), we found that sample replicates and brain tissue types were tightly clustered independent of extraction method (**Figure S11b**). We also found CpGm levels were highly correlated within sample replicates regardless of extraction method and to a much greater degree than between tissue types of the same samples (**Figure S11c**). These results indicated that extraction method (instrument vs manual) does not significantly contribute to differences in DNA methylation levels. Consequently, we performed automated DNA extraction for TCX samples and manual for CER samples.

All study samples were randomized for brain tissue (TCX, CER), sex, age at death, *APOE*ε4 dose and CAA score for sample prep and sequencing. Library preparation and sequencing was performed in the same manner as that for the pilot study. One TCX sample failed sequencing resulting in N= 200 CER and N= 471 TCX samples with available DNAm data. Raw sequencing reads were analyzed using the SAAP-RRBS pipeline described above. Rigorous quality control of the CpG sites was then performed independently for each brain tissue type by 1) removing samples with a bisulfite conversion ratio < 98%, 2) removing sites with less than 10x total read coverage and those that mapped to the mitochondrial chromosome, 3) removing samples that failed genome-wide association study QC performed in a previous study^82^, 4) removing sites with a total read coverage in the 99.9^th^ percentile likely indicative of PCR bias, 5) removing samples that failed sex check by calculating the average X and Y chromosome read coverage normalized to the average read coverage of chromosome 1 defining failed samples as those that are beyond 4 standard deviations of the mean of their respective female and male clusters, 6) removing CpG sites that mapped to the sex chromosomes, 7) masking individual sites for each sample separately that are C to T nucleotide transitions on the forward strand or G to A transitions on the reverse strand as determined from the imputed genotype data^82^, 8) removing outlier samples determined from a principle component analysis (PCA) of the top 50% most variable CpGm sites based on standard deviation using the R function *prcomp*(.scale = TRUE, center = TRUE) with outliers defined as those 4 times the standard deviation from the mean of principal components (PC) 1-3, 9) removing sites not present in over 50% of the samples, and 10) removing samples with less than 50% of the CpGs that passed QC (**Table S2**).

For analyses investigating regional CpGm (rCpGm), we leveraged the ChromHMM core 15-state chromatin model developed by the Roadmap Epigenomics Consortium^48^ for the adult brain Inferior Temporal Lobe (E072). The hg38lift predicted chromatin bed file was downloaded from https://egg2.wustl.edu/roadmap/data/byFileType/chromhmmSegmentations/ChmmModels/coreMarks/jointModel/final/ on June 29, 2022. Although there were regions in this file that were not annotated to a specific chromatin state as a result of the lift over from hg19 to hg38, these individual regions were kept and annotated as “16_NA” or “Not Defined” for downstream analyses. We grouped all CpGm sites that passed QC into regions based on the 15-state chromatin model annotations and averaged each group to determine rCpGm levels (**Figure 1**). rCpGm start and stop sites were defined as the first and last CpGm present in each rCpGm group. As a predicted chromatin state model for the CER did not exist at the time of this study and the fact that we want to analyze rCpGms across tissue types, we used the Inferior Temporal Lobe model to define rCpGms for all brain tissue types. In this study, we had two tiers of DNA methylation measures, one at an individual CpGm site level (CpGm) and one at a regional level according to our novel method defined by the chromatin state model (rCpGm). For each tier, we then performed data reduction steps to remove non-variable sites/regions that are not statistically informative and that would unnecessarily contribute to increasing the multiple testing correction burden. These data reduction steps include, 11) removing CpGm/rCpGm with a methylation ratio range less than 1% as these are likely not biologically informative, and 12) removing sites/regions where 95% or more of the samples are fully methylated (100%) or fully unmethylated (0%) as these provide unreliable statistical estimates (**Table S2**).

### Gene Expression Measures

As part of the MC-CAA study, bulk RNAseq data was also generated for 477 TCX and a subset of 200 CER samples. ***Sample Preparation and Sequencing***: Total RNA was extracted for each tissue sample using Trizol® reagent and cleaned using Qiagen RNeasy columns with DNase treatment. RNA integrity (RIN) was measured using an Agilent Technologies 2100 Bioanalyzer and only samples with a RIN > 5.5 were sent to the Mayo Clinic Genome Analysis Core (GAC) for library preparation and sequencing. The TruSeq RNA Library Prep Kit v2 (Illumina, San Diego, CA) was used for library preparation (non-stranded). Library concentration and size distribution were determined on an Agilent Bioanalyzer DNA 1000 chip. Samples were randomized across sequencing flowcells with respect to average CAA score, sex, age at death, and *APOE*ε4 genotype. The Illumina HiSeq 4000 was used for sequencing with 100bp paired-end reads, multiplexing six samples per flowcell lane. ***Quality Control*:** The Mayo Clinic MAP-Rseq pipeline v.3.0.2^152^ was used for read alignment to hg38 implementing STARv2.5.2b^153^, and featureCounts from the Subread package v1.5.1 for counting^154^. Raw read counts were log2-transformed and normalized using conditional quantile normalization (CQN) via the Bioconductor package; accounting for sequencing depth, gene length, and GC content^155^. FastQC was used for QC of raw sequence reads, and RSeQC^156^ was used for QC of mapped reads. We excluded samples with < 50 million reads mapped to genes, < 50% of reads mapped to genes, and outliers (1.5 inter-quartile range) based on distribution of % junction reads. Sex check was performed by comparing expected expression of Y chromosome genes to recorded sex. Samples that were outliers (>4 standard deviations from the mean) on PCs 1 and 2 in a PCA (*prcomp*, CPM, R statistical software) and multiple other QC variables (RSeQC) were also excluded. We also excluded samples that failed previous genetic based QC steps including relatedness and heterozygosity checks as described previously^82^. There were 456 TCX and 187 CER samples that passed QC and of these, 449 TCX and 184 CER samples had matching RRBS data. We included in our downstream analysis only autosomal genes expressed above background which we defined based on the distribution of CQN values filtering out genes with a CQN value less than 1 for both the CER and TCX datasets.

### AMP-AD datasets

The Mayo RNAseq study^68^, The Mount Sinai Brain Bank (MSBB) study^69^, and The Religious Orders Study and Memory and Aging Project (ROSMAP) Study^70^ were obtained from the AD-knowledge portal (https://adknowledgeportal.synapse.org). The RNA-seq data previously underwent consensus reprocessing by Accelerating Medicines Partnership-AD (AMP-AD, RNAseq Harmonization Study)^71^. Additional QC and diagnosis harmonization of these datasets based on neuropathological measures are described in detail elsewhere^82^.

## Statistical Analyses

### Power Calculation

Power calculations were performed with the *pwr.r.test* function from the *pwr* package v1.3-0 in R (v4.0.2). For a sample size of 472 in a two-sided test, we have 80% power to detect a correlation coefficient of 0.239 and 0.129 at an alpha of 1E-05 and 0.05, respectively. For a sample size of 200 in a two-sided test, we have 80% power to detect a correlation coefficient of 0.359 and 0.197 at an alpha of 1E-05 and 0.05, respectively.

### Phenotypes

Spearman rank correlation between all biochemical measures and CAA have been reported previously^10^ and were consistent with the subset of samples analyzed in this study (data not shown).

### Cell Type Proportion Estimates

Neuronal cell type proportions were estimated from the bulk tissue RNAseq data using the Digital Sorting Algorithm (DSA)^157^ via the *DSA* R package (v1.0). The top 100 genes from BRETIGEA^66^ were used as the cell type specific marker genes for these estimations. Oligodendrocyte precursor cell (OPC) types were not investigated as we previously did not observe robust correlations among the defined OPC marker genes as described in Wang et al 2020^91^.

### Epigenome Wide Association Studies (EWAS)

Primary association models were run using the linear regression *lm*() function in R v4.1.2 adjusting for age at death, sex, and the first three genetic PCs. The CpGm variable (CpGm or rCpGm) was included in each association model as the dependent variable. There were three samples with a Thal phase measure of 2 (2 TCX and 1 with both TCX and CER) that passed DNAm QC. Given the low number of this measure in a single group and the possibility that small variations in this group may considerably and unreliably impact our statistical estimates, we set these Thal phase 2 measures to *NA* in analyses that included Thal phase as a variable. False Discovery Rate (FDR) corrected p-values were calculated with the R *p.adjust* package using the Benjamini & Hochberg method.

### Association with Gene Expression measures

In the MC-CAA dataset, there were N= 449 TCX and N= 184 CER samples that had both RRBS and bulk RNA seq data that could be investigated. *Cis-*rCpGm-Gene pairs were identified by connecting rCpGm with all genes +/- 500kb away from the start and end of each rCpGm for the TCX and CER separately. Residuals of rCpGm were extracted by fitting the linear regression model: rCpGm ∼ AgeAtDeath + Sex + genetic PC 1-3. Bulk gene expression (GE) residuals were extracted by fitting the mixed effects linear regression model: GE ∼ AgeAtDeath + Sex + RIN + (1|batch). The association between rCpGm and bulk gene expression residuals were tested by fitting the linear regression model: Gene_Expression_residuals ∼ rCpGm_residuals for each *cis-*rCpGm-Gene pair independently for each tissue type.

To identify gene expression associations with the AD endophenotypes, we tested the association of normalized gene expression data with the endophenotype of interest. For continuous phenotypes (biochemical measures and CAA), we ran the following linear regression model: cqn(gene expression) ∼ ADphenoytpe + Age at Death + sex + RIN + (1|flowcell) using the *lmer* function in the *lme4* package^158^. For ordinal variables (Braak stage and Thal phase), we first extracted residuals from the gene expression values using the following model: cqn(gene expression) ∼ Age at Death + sex + RIN + (1|flowcell), and then associated the gene expression residuals with each phenotype in an ordinal regression model using the *clm* function in the *ordinal* R package. As with the DNAm data, because there were only a small number of samples with a Thal phase of 2, these were set to *NA* in analyses that included Thal as a variable. FDR p-values were calculated with the R *p.adjust* package using the Benjamini & Hochberg method.

The AMP-AD datasets were used to investigate the association between CQN normalized gene expression levels of implicated genes with Braak stage in non-Hispanic white, neuropathologically defined AD cases and controls using linear regression in R with the *lm()* function. Mayo TCX models adjusted for age at death, sex, RNA integrity number (RIN), and sequencing flow cell rounding down intermediate Braak stage numbers to the next lowest stage. MSBB BM22 models adjusted for age at death, sex, RIN, and batch. ROSMAP models adjusted for age at death, sex, RIN, sequencing batch, and study (ROS/MAP). Individuals with an age at death above 90 were set to 90, in compliance with HIPAA rules. Oligodendrocyte proportions were further adjusted for in each model by including the *OLIG2* (ENSG00000205927) normalized expression levels as a covariate.

### mQTL analysis

*Cis-*methylation Quantitative Trait Locus (*cis-*mQTL) analysis was performed using the R package *MatrixEQTL*^159^ for all genetic variant dosage and rCpGms pairs +/- 500kb apart for the TCX and CER separately adjusting for age at death, sex, and the first 3 genetic PCs. In total, we investigated 289,078,059 TCX rCpGm-SNP pairs and 291,400,204 CER rCpGm-SNP pairs. False Discovery Rate (FDR) corrected p-values were calculated with the R *p.adjust* package using the Benjamini & Hochberg method.

### Sensitivity Analyses

We performed post-hoc sensitivity analyses to test the robustness of the top associations to effects of batch, neuropathological changes, and neuronal cell type proportions. To determine if top associations were influenced by batch, we included batch in a linear regression model as a random effect variable using the *lmer()* function of the *lme4* R package^158^: rCpGm ∼ Endophenotype + Age at death + Sex + Genetic PCs 1-3 + (1|FlowCell). To determine if top biochemical associations were influenced by neuropathological changes, we included Braak stage and Thal phase as fixed-effect covariates in a linear regression model: rCpGm ∼ Endophenotype + Age at death + Sex + Genetic PCs 1-3 + Braak stage + Thal phase. To determine if top associations were influenced by neuronal cell type proportions, we included DSA estimated neuronal cell type proportions as a fixed-effect covariate in the linear regression model: rCpGm ∼ Endophenotype + Age at death + Sex + Genetic PCs 1-3 + Neuronal Cell Proportion. Lastly, to determine if distribution of a phenotype influenced the top associations, we performed Spearman rank correlation analysis in R using the *cor.test()* function of the *stats* package.

### Pathway analysis

Pathway enrichment was tested using the Gene Ontology Biological Processes database in the *anRichment* R package with p-values computed via a one-sided hypergeometric test.

### Annotations

CpGm sites were annotated (Island, shore, shelf, inter-genic) with the R package *annotatr()* v1.21.0^160^ querying AnnotationHub_3.2.2 in R (v4.1.2).

### DNAm Replication Datasets

Data from The Religious Orders Study and Memory and Aging Project (ROSMAP) Study was downloaded from the AD knowledge portal for 707 Non-Hispanic White, dorsolateral prefrontal cortex (DLPFC) samples with DNAm data measured on the Illumina HumanMethylation450 BeadChip array^30^ including IDAT and metadata files on May 17^th^, 2023 (**Table S3**). Additional and updated metadata for these ROSMAP samples was also obtained from the Rush Alzheimer’s Disease Center (RADC) on May 22^nd^, 2023. Data from the Brains for Dementia Research (BDR) Dataset was downloaded for 630 individuals with DLPFC and occipital cortex (OCC) DNAm measured on the Illumina EPICv1 DNA methylation array^32^ including IDAT files from the Gene Expression Omnibus (GEO, accession number GSE197305) on February 23^rd^, 2023 and metadata from the UK Brain Bank Network (UKBBN) database after project approval (https://brainbanknetwork.ac.uk/) on May 26^th^, 2023 (**Table S3**). PSP data from Weber *et al.* 2018^73^ was downloaded from GEO (GSE75704) on April 12^th^, 2023 for 94 PSP and 72 control individuals with prefrontal lobe DNAm measured using the Illumina HumanMethylation450 BeadChip. The ROSMAP, BDR, and Weber PSP datasets were consensus reprocessed through a unified QC pipeline for each brain tissue type separately using R (v4.1.2). For consistency with the Mayo Clinic discovery cohort, individuals with an indicated ethnicity other than Non-Hispanic white or white were removed, individuals with unknown ethnicity were kept. Raw IDAT files were imported as a RGChannelSet object using *read.metharray.exp* in *minfi*^161^. Sample and probe QC was performed using the following steps: 1) excluding probes that mapped to non-autosomal chromosomes; 2) excluding samples with a low (<80%) bisulfite conversion rate estimated by *bscon* in *wateRmelon*^162^; 3) using detection p-values obtained by *detectionP* in *minfi* to exclude probes where <99% of the samples had a detection p-value>0.05; 4) removing samples where <99% of the probes had a detection p-value >0.05; 5) removing samples that failed sex check by clustering samples on the first PCs of previously identified robust sex-related probes on the X and Y chromosomes as implemented by *estimateSex*^163^ in *wateRmelon*; 6) checking genotype consistency and removing mismatched samples by comparing the β-value means from the SNP probes for individuals with samples from multiple brain regions; 7) removing probes that have previously been identified to have cross-hybridizing potential or polymorphic targets (with European MAF>0.01)^164^, as well as the SNP and control probes; 8) removing outlier samples defined by the *outlyx* function in *wateRmelon*. Subsequently, to minimize intra- and inter-array variation, as well as probe design (I/II) bias, quantile normalization was performed using *dasen*^162^ in *wateRmelon*, and outlier check was performed again on the normalized data. Individual sex and age at death were compared when available from different sources and when discrepancies were identified, individuals were removed from the analysis. PCA was performed with the normalized β-values using *prcomp*(.*scale* = TRUE, *center*= TRUE) on the top 50% most variable CpGm sites based on standard deviation. Non-CpGm sites were removed and CpGm sites were lifted to the hg38 genome using the *liftOver* package and corresponding chain files provided by UCSC Genome Browser. rCpGms were calculated and data reduction steps were performed as described for the RRBS data. Intermediate Braak stages were grouped with the next lowest stage. AD cases were neuropathologically defined as individuals with a Braak stage ≥ 4 and a Thal phase ≥ 2. Control samples were defined as individuals with a Braak stage ≤ 4 and a Thal phase ≤ 2 (**Table S3**). Any donor who was not neuropathologically defined as an AD case or control was designated as “Other”. For the Weber PSP dataset, PSP and control sample definitions were used as described^73^. For ROSMAP and BDR, EWAS were performed in each dataset and tissue type separately for Braak stage, Thal phase, and AD diagnosis looking at AD vs Controls using the *lmer* function in the *lmerTest* R package. For the ROSMAP linear mixed models, we adjusted for age at death, sex, study (ROS/MAP), and batch as fixed effects as well as slide as a random effect variable. For the BDR linear mixed models, we adjusted for age at death, sex, and CpGm PC’s 1-3 as fixed effects as well as slide as a random effect variable. We included CpGm PC’s 1-3 in the BDR models because batch, although not available, has previously been found to be a source of variation in this dataset and captured by PC’s^32^. PC’s 1-3 were chosen because these explained more than 5% of variation (PC1-23%, PC2-13%, PC3-6%) cumulating to a total of 41%. For Weber PSP, EWAS were run using *lmer* and linear mixed models for PSP diagnosis and Braak stage adjusting for age at death, sex, and CpGm PC’s 1-3 as fixed effects as well as slide as a random effect variable. Similarly, because we did not have information on batch, we included PC’s 1-3 explaining 25%, 12%, and 5% of the variation, respectively. Next, due to consistently high level of genomic inflation, we used the *bacon* R package^165^ to adjust for inflation in each of the EWAS models (**Table S18**).

### Previously published single nuclei/cell RNAseq studies

Oligodendrocyte specific gene expression association results from previously published single nuclei/cell RNAseq studies depicted in **Figure 5B** and **Figure S10** were downloaded in October, 2024 from the following publications: Is 2024^74^, Mathys 2023^75^, Gamache 2023^76^, Morabito 2021^77^, Anderson 2023^78^, Rexach 2024^79^, Min 2023^80^.

### Cell type marker gene enrichment

One sided hypergeometric tests using the *phyper(lower.tail = FALSE)* function from the R *stats* package were used to test for enrichment of the top 100 and top 1000 cell type marker genes identified by BRETIGEA^66^ in our dataset. We used the total number of genes expressed above background tested in our gene expression analysis (N= 18,383 genes) as the total universe number.

## Acknowledgements

We thank the patients and families for their participation and tissue donation, without whom these studies would not have been possible. We thank Özkan İş, Jianna Tan, and Jeremiah Bergman for their technical assistance. We thank the Mayo Clinic Genome Analysis Core (GAC), Co-Directors, Julie M. Cunningham, PhD and Eric Wieben, PhD, and supervisor Julie Lau, for their collaboration in collection of omics data. We also thank our colleague Saurabh Baheti at the Mayo Clinic Bioinformatic Core (BIC) for their collaboration in data quality control.

**AD Knowledge Portal:** The results published here are in whole or in part based on data obtained from the AD Knowledge Portal (https://adknowledgeportal.org). Mayo Clinic AD-CAA: The Mayo Clinic AD-CAA study was led by Dr. Guojun Bu and Dr. Nilüfer Ertekin-Taner at the, Mayo Clinic, Jacksonville, FL as part of the multi-PI RF1AG051504 (MPIs Bu and Ertekin-Taner) using samples from the Mayo Clinic Brain Bank. Data collection was supported through funding by NIA grants P50AG016574, R37AG027924, Cure Alzheimer’s Fund, and support from Mayo Foundation. <u>AMP-AD datasets</u>: The results published here are in whole or in part based on data obtained from the AMP-AD Knowledge Portal (https://doi.org/10.7303/syn2580853). Mayo Clinic: The Mayo RNAseq study data was led by Dr. Nilüfer Ertekin-Taner, Mayo Clinic, Jacksonville, FL as part of the multi-PI U01 AG046139 (MPIs Golde, Ertekin-Taner, Younkin, Price). Samples were provided from the following sources: The Mayo Clinic Brain Bank and Banner Sun Health Research Institute. Data collection was supported through funding by NIA grants P50 AG016574, R01 AG032990, U01 AG046139, R01 AG018023, U01 AG006576, U01 AG006786, R01 AG025711, R01 AG017216, R01 AG003949, NINDS grant R01 NS080820, CurePSP Foundation, and support from Mayo Foundation. Study data includes samples collected through the Sun Health Research Institute Brain and Body Donation Program of Sun City, Arizona. The Brain and Body Donation Program is supported by the National Institute of Neurological Disorders and Stroke (U24 NS072026 National Brain and Tissue Resource for Parkinsons Disease and Related Disorders), the National Institute on Aging (P30 AG19610 Arizona Alzheimers Disease Core Center), the Arizona Department of Health Services (contract 211002, Arizona Alzheimers Research Center), the Arizona Biomedical Research Commission (contracts 4001, 0011, 05-901 and 1001 to the Arizona Parkinson’s Disease Consortium) and the Michael J. Fox Foundation for Parkinsons Research. MSBB: These data were generated from postmortem brain tissue collected through the Mount Sinai VA Medical Center Brain Bank and were provided by Dr. Eric Schadt from Mount Sinai School of Medicine. ROSMAP: Study data were provided by the Rush Alzheimer’s Disease Center, Rush University Medical Center, Chicago. Data collection was supported through funding by NIA grants P30AG10161 (ROS), R01AG15819 (ROSMAP; genomics and RNAseq), R01AG17917 (MAP), R01AG30146, R01AG36042 (5hC methylation, ATACseq), RC2AG036547 (H3K9Ac), R01AG36836 (RNAseq), R01AG48015 (monocyte RNAseq) RF1AG57473 (single nucleus RNAseq), U01AG32984 (genomic and whole exome sequencing), U01AG46152 (ROSMAP AMP-AD, targeted proteomics), U01AG46161(TMT proteomics), U01AG61356 (whole genome sequencing, targeted proteomics, ROSMAP AMP-AD), the Illinois Department of Public Health (ROSMAP), and the Translational Genomics Research Institute (genomic). Additional phenotypic data can be requested at www.radc.rush.edu. AGORA: The results published here are in whole or in part based on data obtained from Agora, a platform initially developed by the NIA-funded AMP-AD consortium that shares evidence in support of AD target discovery (agora.adknowledgeportal.org/). RADC: We thank the study participants and staff of the Rush Alzheimer’s Disease Center. ROSMAP^166^ is supported by P30AG10161, P30AG72975, R01AG15819, R01AG17917, U01AG46152, and U01AG61356. ROSMAP resources can be requested at https://www.radc.rush.edu and www.synpase.org. All ROSMAP participants are enrolled without known dementia and agreed to detailed clinical evaluation and brain donation at death. Both ROS and MAP studies were approved by an Institutional Review Board of Rush University Medical Center. Each participant signed an informed consent, Anatomic Gift Act, and an RADC Repository consent allowing their data and biospecimens to be repurposed.

UK Brain Bank Network: Brains for Dementia Research: The provision of data used in this study was supported by the UK Brain Banks Network with funding from the UK Medical Research Council and BDR (Brains for Dementia Research).

## Author Contributions

SRO and NET wrote the manuscript; SRO, NET, and MA designed the study; SRO, ZQ, AA, JSR, XW, MMC and FV performed data analyses; MH consulted on statistical methods; SRO built the web application; YM assisted with web application development; CCL, YY, YAM, NZ, TK, and GB collected biochemical measures; DWD, MD, and MEM provided neuropathological data and tissue samples; TTN and KM isolated DNA from tissue samples; NET oversaw the study and provided direction, funding, and resources. All authors reviewed and contributed to the manuscript. The authors read and approved the final manuscript.

## Abbreviations

A1: Tested Allele
Aβ: Amyloid-β
AD: Alzheimer’s disease
AMP-AD: Accelerating Medicines Partnership-Alzheimer’s Disease
APOE: apolipoprotein E
APP: Amyloid Precursor Protein
β: Beta value
BDR: Brains for Dementia Research
BIC: Bioinformatics core
BP: Base Pair
CAA: Cerebral amyloid angiopathy
CER: Cerebellum
CI: Confidence Interval
CpGm: DNA methylation at a CpG site
CQN: Conditional Quantile Normalization
DLPFC: Dorsal lateral prefrontal cortex
DNAm: DNA methylation
DSA: Digital Sorting Algorithm
EWAS: Epigenome-wide association study
FA: Formic acid soluble tissue fraction
FDR: False Discovery Rate
GAC: Genome analysis core
GE: Gene expression
GEO: Gene Expression Omnibus
GWG: Genome-wide genotypes
HRC: Haplotype reference consortium
ln: natural log
MAF: Minor Allele Frequency
MC-CAA: Mayo Clinic CAA study
mQTL: methylation quantitative trait locus
MSBB: Mount Sinai Brain Bank
N: Number
NFT: Neurofibrillary tangles
NIA: National Institutes of Aging
OCC: Occipital Cortex
P: P-value
PC: Principal Component
PCA: Principal Component Analysis
PCR: Polymerase chain reaction
PSP: Progressive Supranuclear Palsy
pTau: phosphorylated tau
QC: Quality Control
RADC: Rush Alzheimer’s Disease Center
rCpGm: DNA methylation across a region of CpG sites (regional CpGm)
ROSMAP: Religious Orders Study and Rush Memory Aging Project
RRBS: Reduced Representation Bisulfite Sequencing
SAAP-RRBS: Streamlined Analysis and Annotation Pipeline for Reduced Representation Bisulfite Sequencing
scRNA-seq: Single cell RNA sequencing
SD: Standard Deviation
SE: Standard Error
Sqrt: Square root
sqrt(CAA): Square root transformed CAA scores
TBS: Tris-buffered saline soluble tissue fraction
TCX: Temporal cortex
TSS: Transcription Start Site
TssA: Active TSS
TX: Detergent (1% Triton-X) soluble tissue fraction
UKBBN: UK Brain Bank Network
WGBS: Whole genome bisulfite sequencing

