## Supplemental Figures for "Integrative Epigenomic Landscape of Alzheimer’s Disease Brains Reveals Oligodendrocyte Molecular Perturbations Associated with Tau"

**Author Affiliations:**

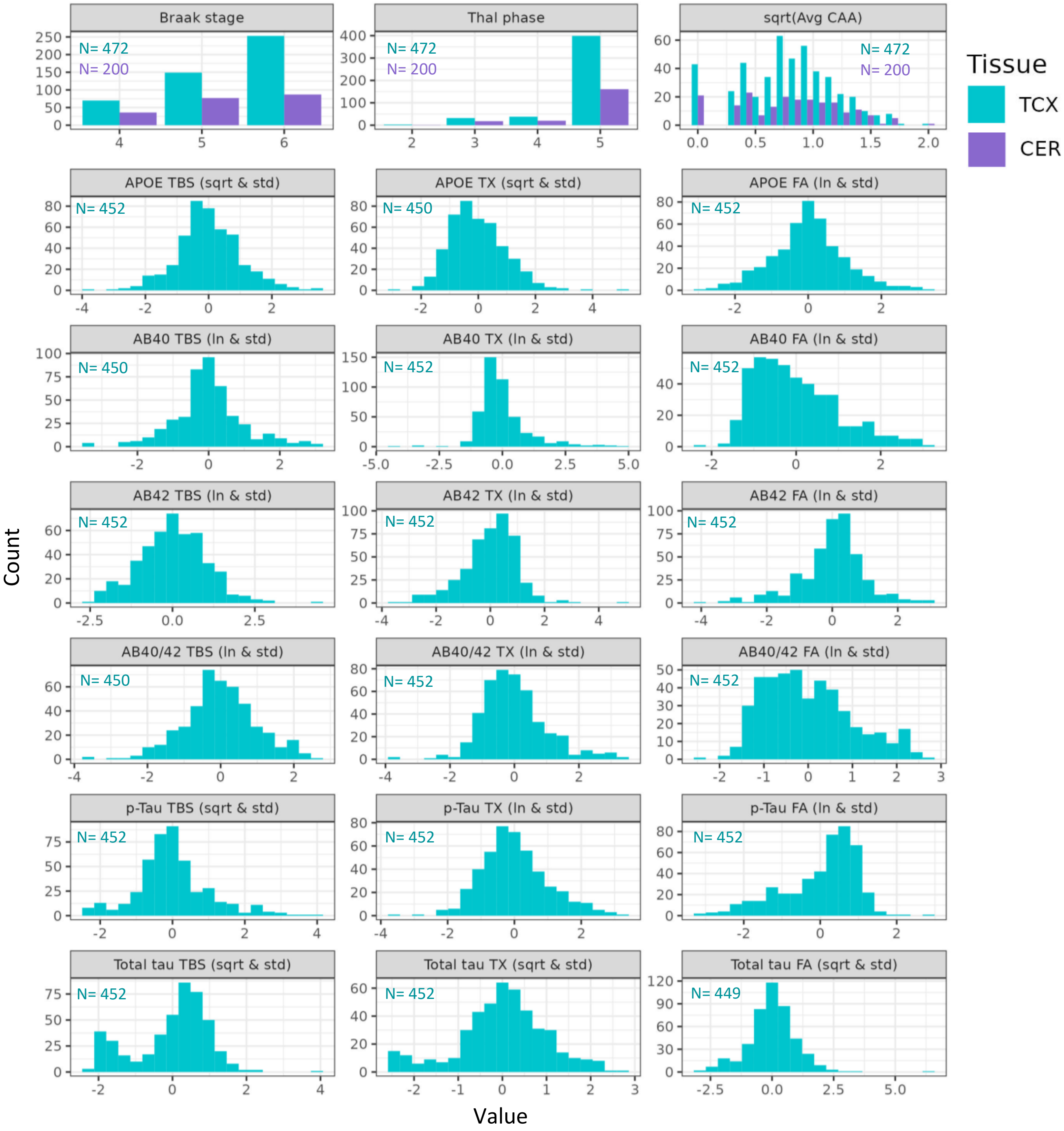


**Figure S1: Histograms of AD endophenotypes measured in each brain tissue investigated**. Histogram plots of the AD endophenotypes of interest for each brain tissue region. Teal color represents the TCX samples and purple the CER samples. Numbers in the upper left corner (N) indicate the number of samples with DNAm and endophenotype measurements for each brain region. Biochemical measures were either natural log (ln) or square root transformed (sqrt) to approximate a normal distribution and then standardized (std). Neuropathology endophenotypes were available for both CER and TCX, whereas biochemical measures were obtained only for TCX samples.

**CER** = Cerebellum, **ln** = natural log transformed, **sqrt** = square root transformed, **std** = standardized, **TCX** = Temporal Cortex

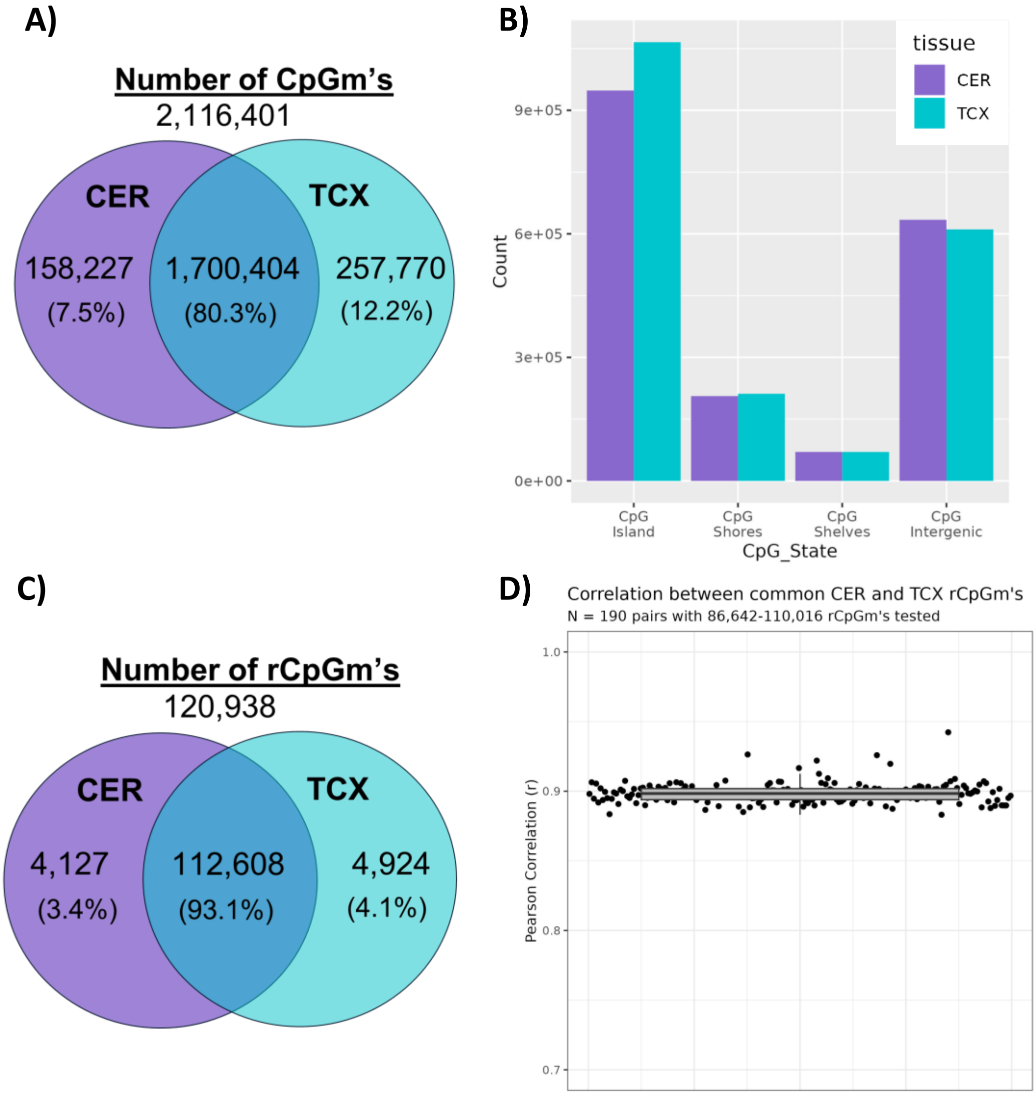


**Figure S2: Summary data of measured DNAm**. A) Venn diagram of individual CpGm sites that passed quality control (QC) between the CER and TCX. B) Histogram of CpG annotations for the individual CpG sites in the CER and TCX. C) Venn diagram of rCpGms that passed QC between the CER and TCX. D) Pearson Correlation estimates between common rCpGm in the CER and TCX in each sample with DNAm from both tissue types (N=190). All correlations had a pvalue < 2.2E-16.

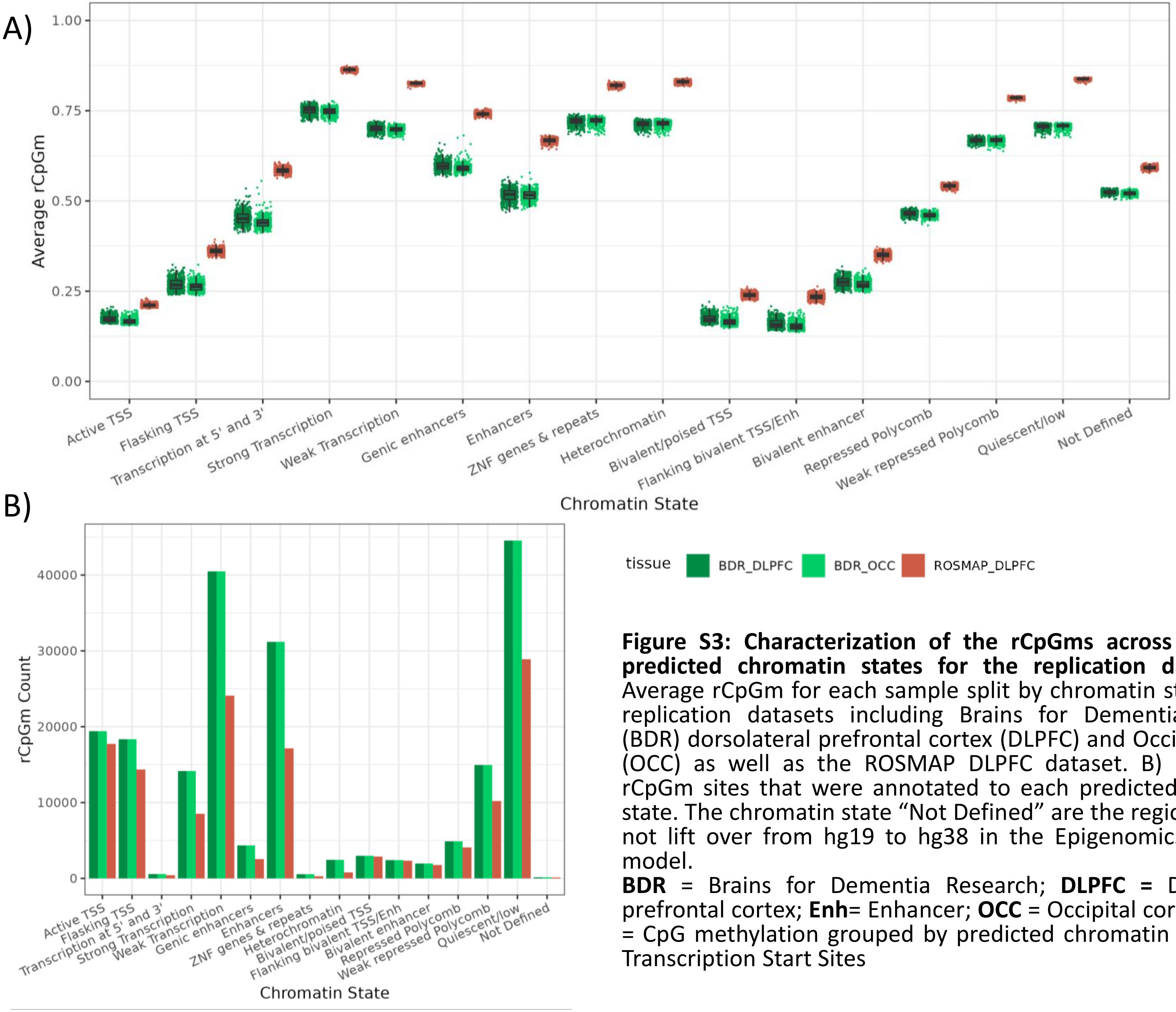


**Figure S3: Characterization of the rCpGms across annotated predicted chromatin states for the replication datasets**. A) Average rCpGm for each sample split by chromatin state for the replication datasets including Brains for Dementia Research (BDR) dorsolateral prefrontal cortex (DLPFC) and Occipital cortex (OCC) as well as the ROSMAP DLPFC dataset. B) Number of rCpGm sites that were annotated to each predicted chromatin state. The chromatin state "Not Defined" are the regions that did not lift over from hg19 to hg38 in the Epigenomics Roadmap model.

**BDR** = Brains for Dementia Research; **DLPFC =** Dorsolateral prefrontal cortex; **Enh**= Enhancer; **OCC** = Occipital cortex; **rCpGm** = CpG methylation grouped by predicted chromatin state; **TSS**= Transcription Start Sites

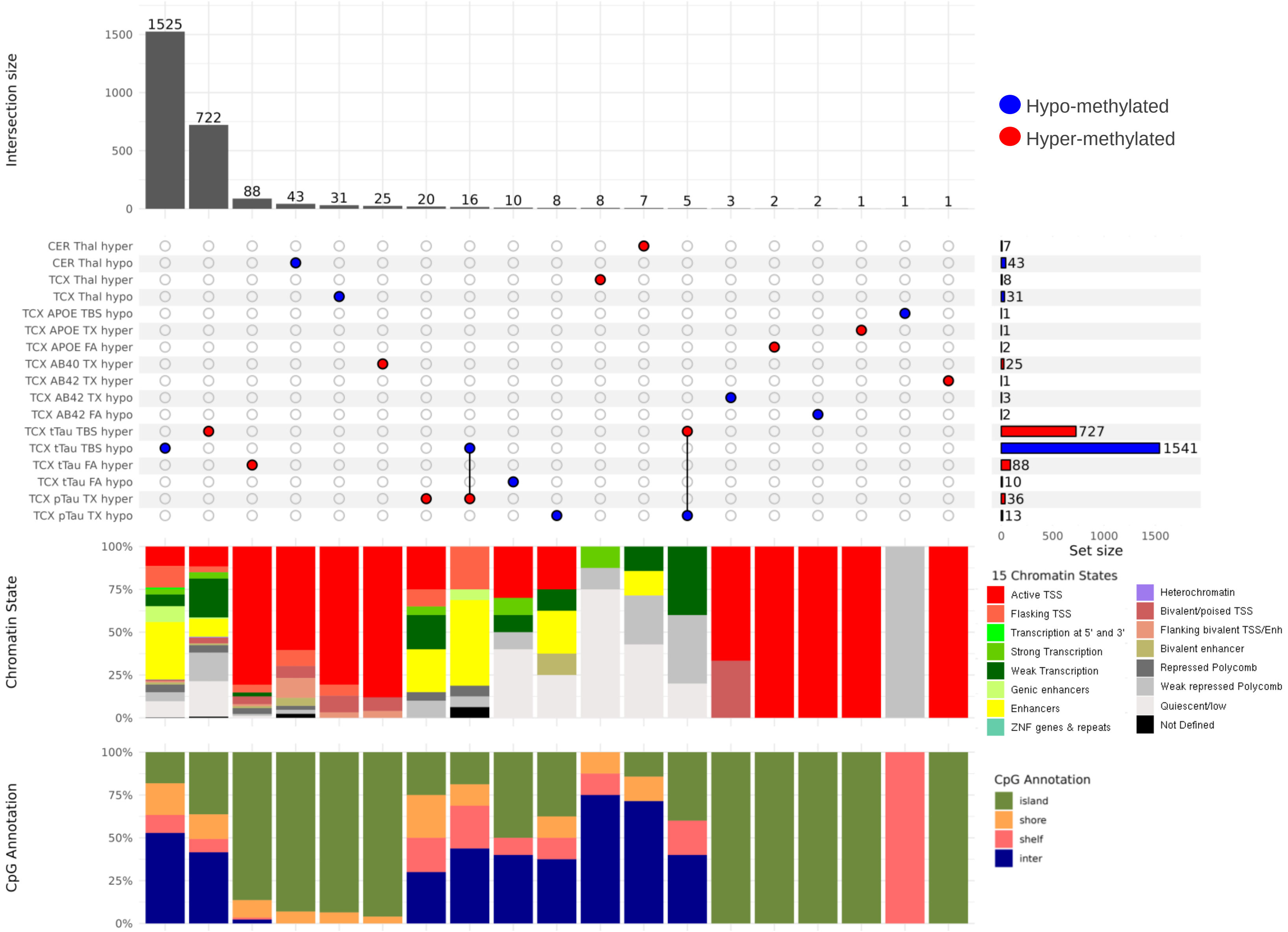


**Figure S4: Upset Plot for FDR Significant Individual CpGm EWAS Results.**
Upset plot showing FDR significant (FDR p-value ≤ 0.05) individual CpGm associations tested in the cerebellum with the neuropathological measures and in the temporal cortex with the neuropathological and biochemical measures. Bar graphs show additional annotations for each intersection set of the FDR significant CpGm associations. Chromatin states depict the Epigenetic Roadmap 15-chromatin state model annotation breakdown for each intersection set. CpG Annotations depict the CpG annotation breakdown for each intersection set. Red indicates hyper-methylation or a positive direction of association and blue indicates hypo-methylation or a negative direction of association with each endophenotype.
**CER** = Cerebellum; **FA** = Insoluble formic acid fraction; **FDR** = false discovery rate; **N** = Number; **pTau** = phospho-tau; **TBS** = soluble tris buffered saline fraction; **tTau** = total tau; **TCX**= Temporal Cortex; **TX**= membrane associated Triton-X fraction

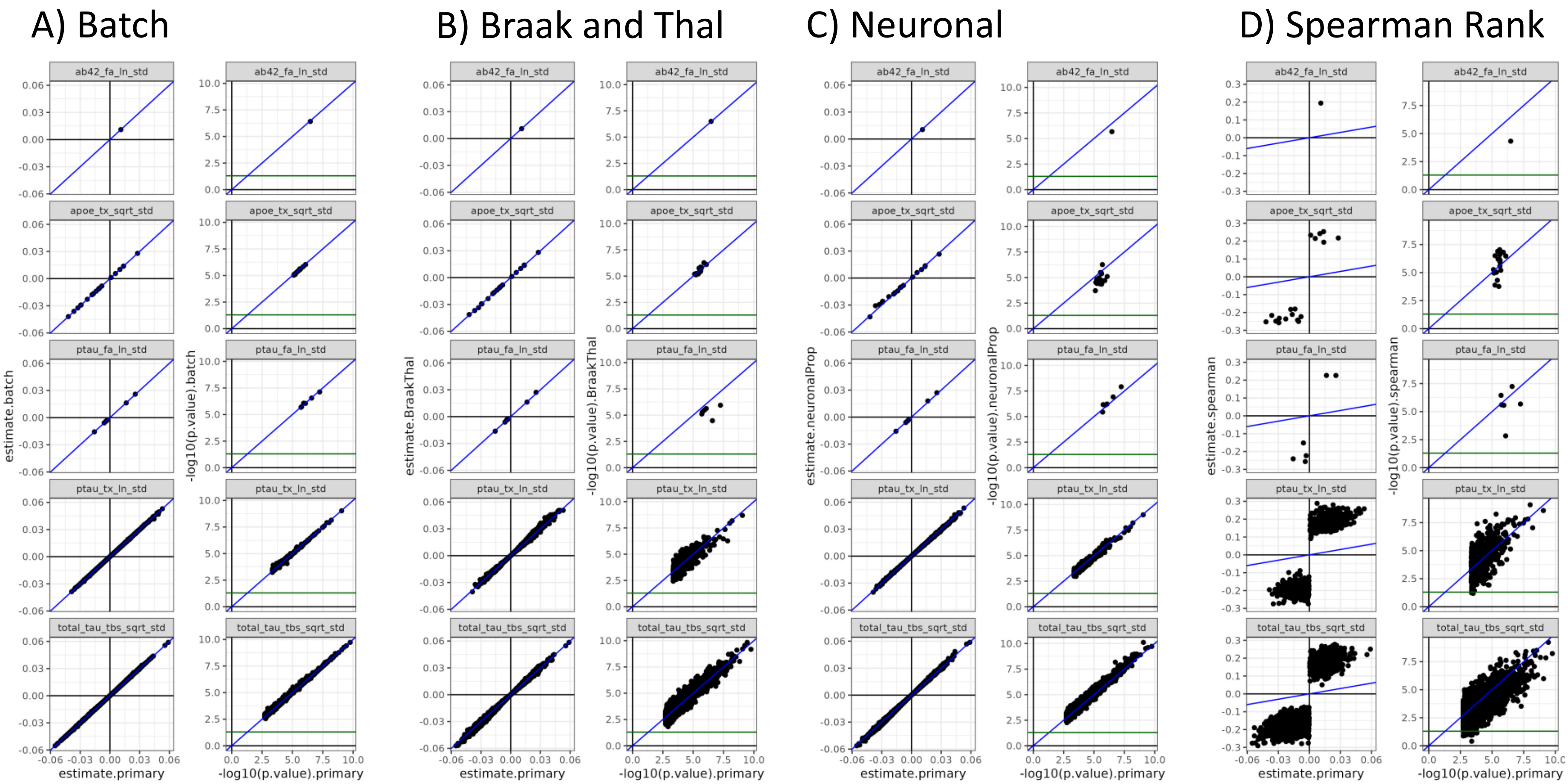


**Figure S5: Comparison of Primary EWAS model results with sensitivity EWAS model results for FDR significant TCX rCpGms**. Comparison of test statistics (estimates and p-values) for all FDR significant rCpGms from the primary rCpGm EWAS models and the sensitivity EWAS models. Sensitivity models included additional covariates in the linear regression models including A) sequencing batch as a random effect, B) Braak stage and Thal phase, C) Neuronal proportion estimated from bulk RNAseq data using DSA, and D) a spearman rank test. Primary model statistics are the x-axes, sensitivity model statistics are the y- axes. Blue diagonal line is the [1,1] line. Green line is the –log10(pval = 0.05) value. **FA**= insoluble tissue fraction, **ln** = natural log transformed, **sqrt** = square root transformed, **std** = standardized, **TBS** = soluble tissue fraction, **TCX** = Temporal Cortex, **TX**= membrane tissue fraction

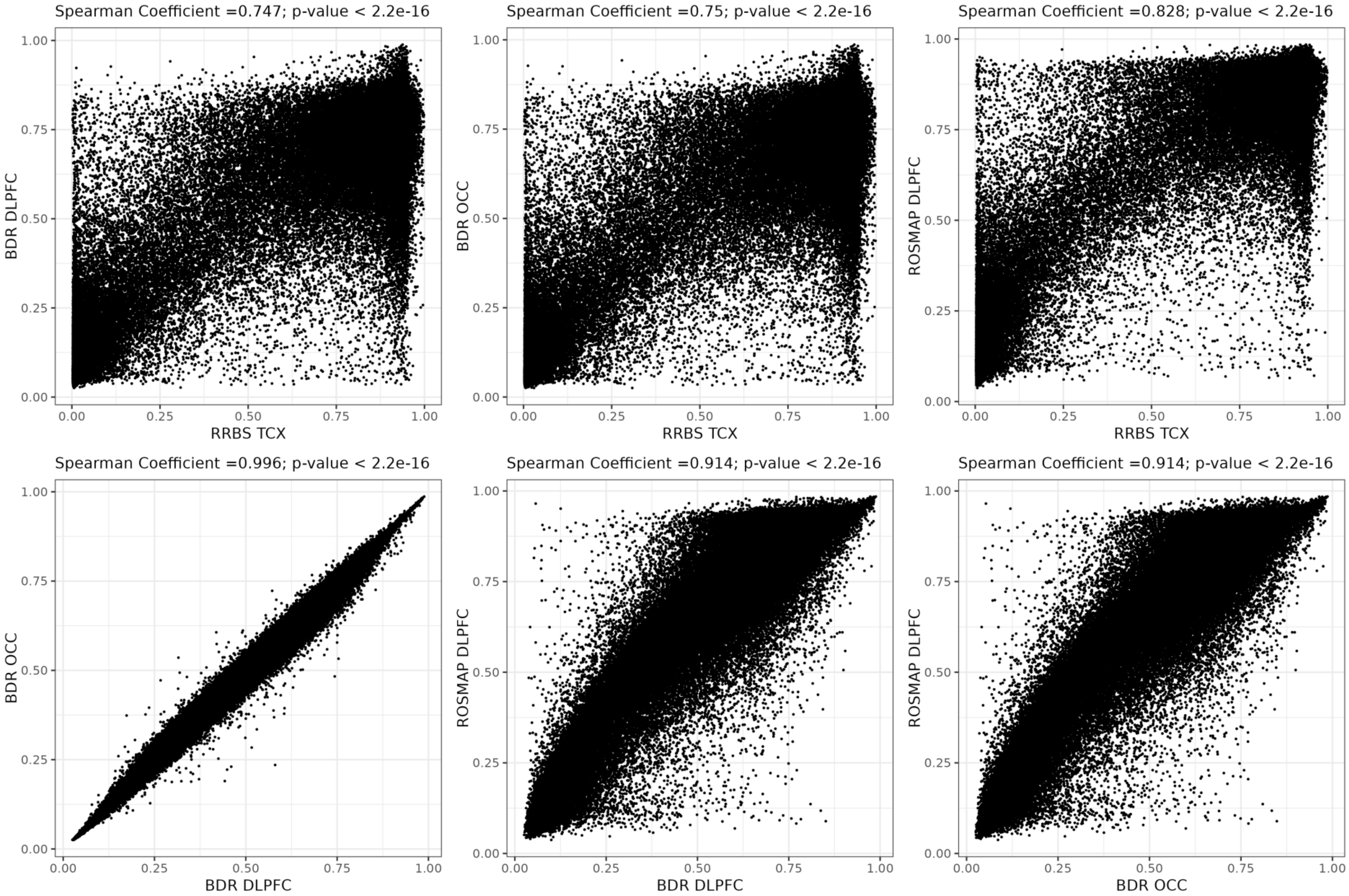


**Figure S6: Correlation of rCpGm levels averaged within a study dataset across samples to overlapping rCpGm sites from other study datasets.** We averaged the methylation values across all samples for each rCpGm in a dataset and then tested the correlation of overlapping rCpGm levels across each dataset via Spearman Correlation. Each point is a corresponding rCpGm pair from each dataset which maps to the same chromatin state window.

**DLPFC** = Dorsolateral Prefrontal Cortex, **OCC** = Occipital Cortex, **TCX** = Temporal Cortex

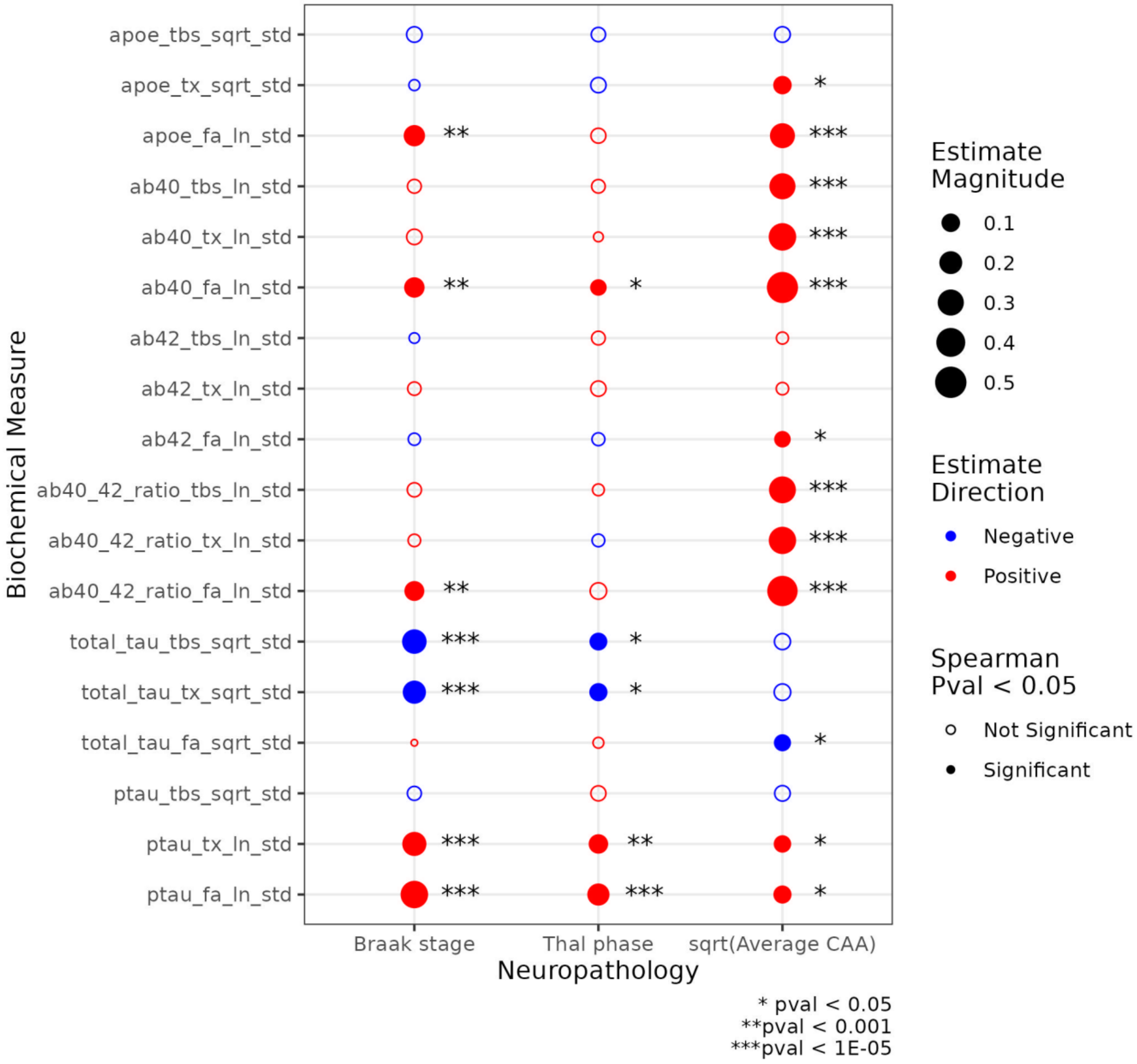


**Figure S7: Spearman Correlation of TCX Biochemical and Neuropathology Measures**.
Two-sided spearman correlation tests were run for each TCX biochemical measure and neuropathology score across all individuals in our Mayo Clinic TCX RRBS study cohort. Sqrt(Average CAA) correlations with each non-standardized biochemical measure were originally published in Liu *et al* 2020. Red indicates a positive direction of correlation, blue indicates a negative direction of correlation, the circle type indicates significance of correlation (filled circle if pvalue < 0.05), absolute value of the correlation coefficient estimate is indicated by dot size.
**FA**= insoluble tissue fraction, **ln** = natural log transformed, **sqrt** = square root transformed, **std** = standardized, **TBS** = soluble tissue fraction, **TCX** = Temporal Cortex, **TX**= membrane tissue fraction

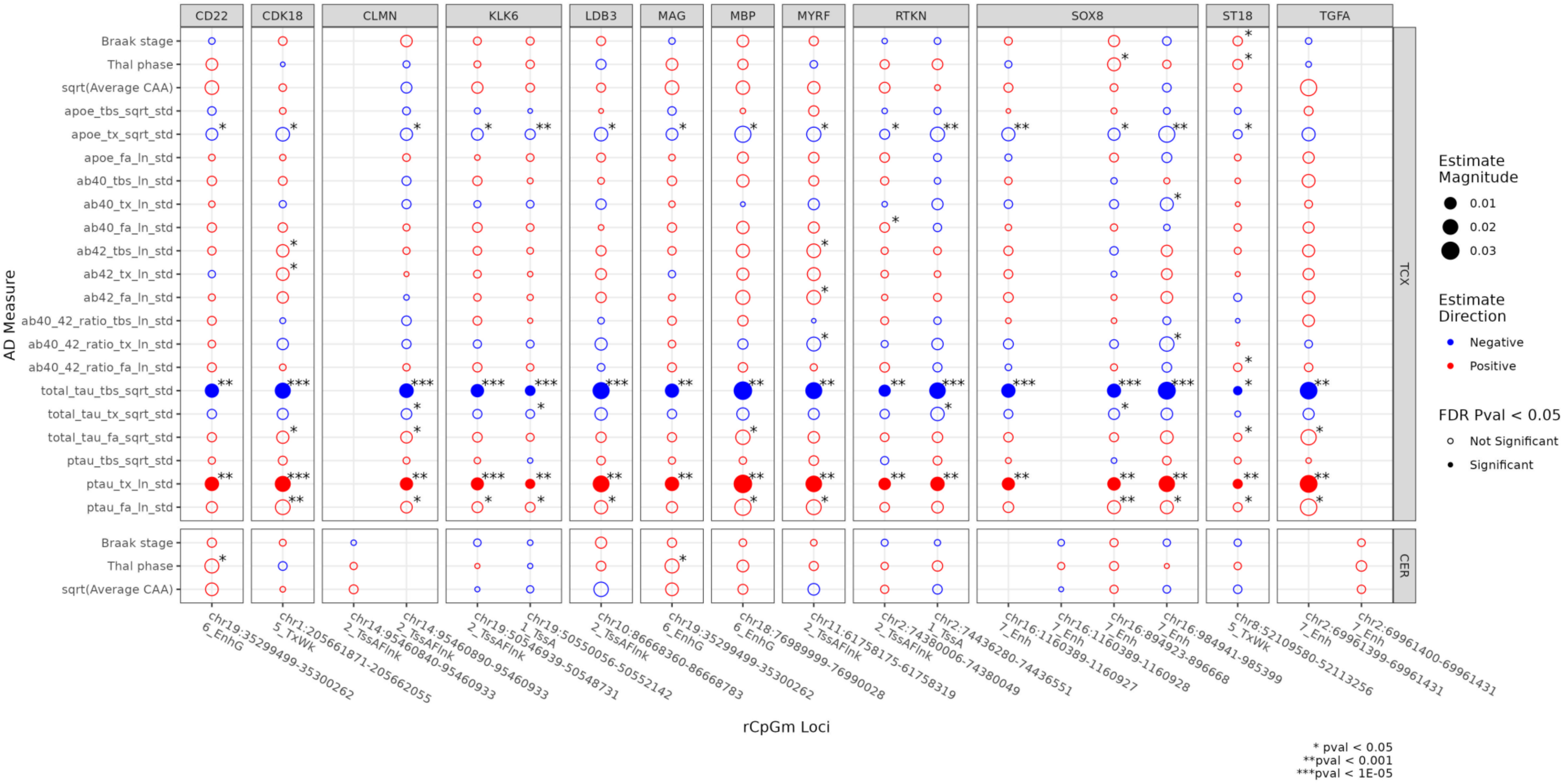


**Figure S8: EWAS results across AD endophenotypes for top oligodendrocyte marker gene rCpGms**.

EWAS association results for the top rCpGms that associate with oligodendrocyte marker gene brain levels. Red indicates a positive direction of association, blue indicates a negative direction of association, the circle type indicates if the is significant at FDR pvalue < 0.05 (solid circle) or not (open circle), absolute value of the estimate is indicated by dot size, asterisk specifies different p-value of associations shown below the figure.

**FA**= insoluble tissue fraction, **ln** = natural log transformed, **sqrt** = square root transformed, **std** = standardized, **TBS** = soluble tissue fraction, **TCX** = Temporal Cortex, **TX**= membrane tissue fraction

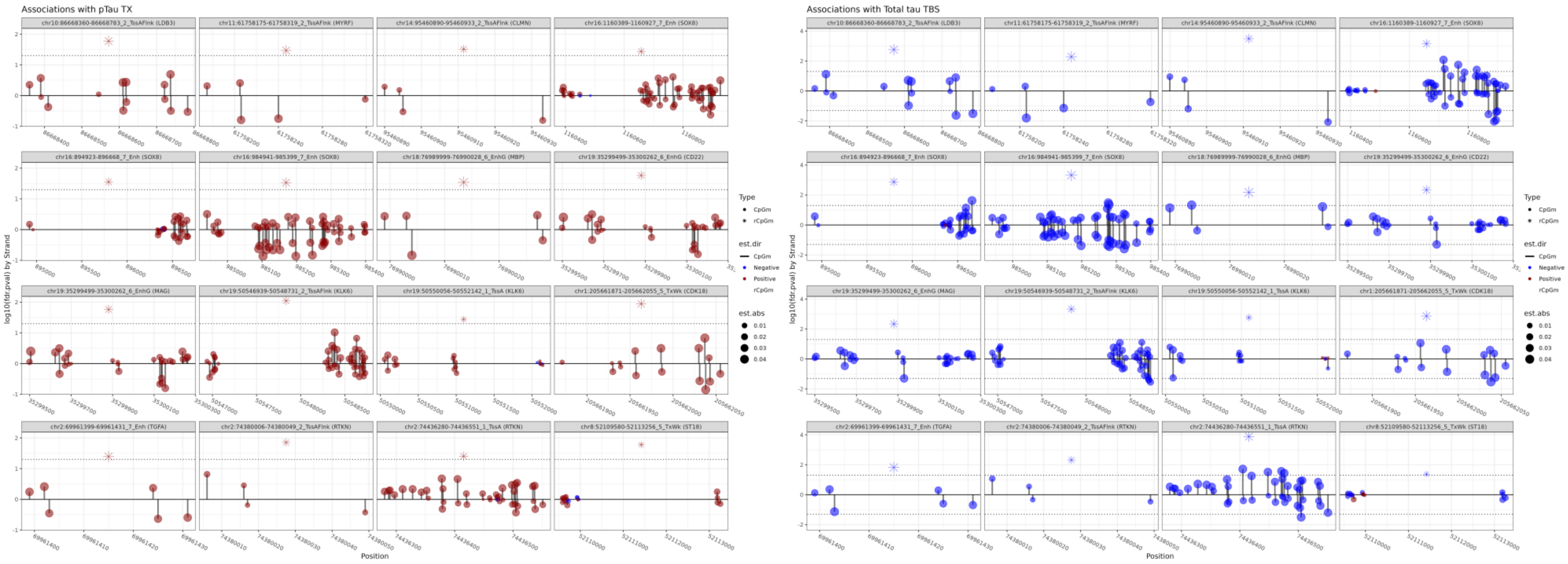


**Figure S9: EWAS results for the individual CpGms comprising the top oligodendrocyte marker gene rCpGms associated with Total tau$_{TBS}$ and pTau$_{TX}$.** Each circle represents an individual CpGm that makes up its corresponding rCpGm represented by a star. Direction of association is indicated by color with a positive association shown as red and negative as blue with size of the circle corresponding to the magnitude of effect. Y-axis indicates log10(FDR pvalue) with positive values indicating the CpG is on the forward strand and negative values on the reverse strand. X-axis is the base pair position of the CpGms while the rCpGms are set to the middle of the region.

**FA**= insoluble tissue fraction, **ln** = natural log transformed, **sqrt** = square root transformed, **std** = standardized, **TBS** = soluble tissue fraction, **TCX** = Temporal Cortex, **TX**= membrane tissue fraction

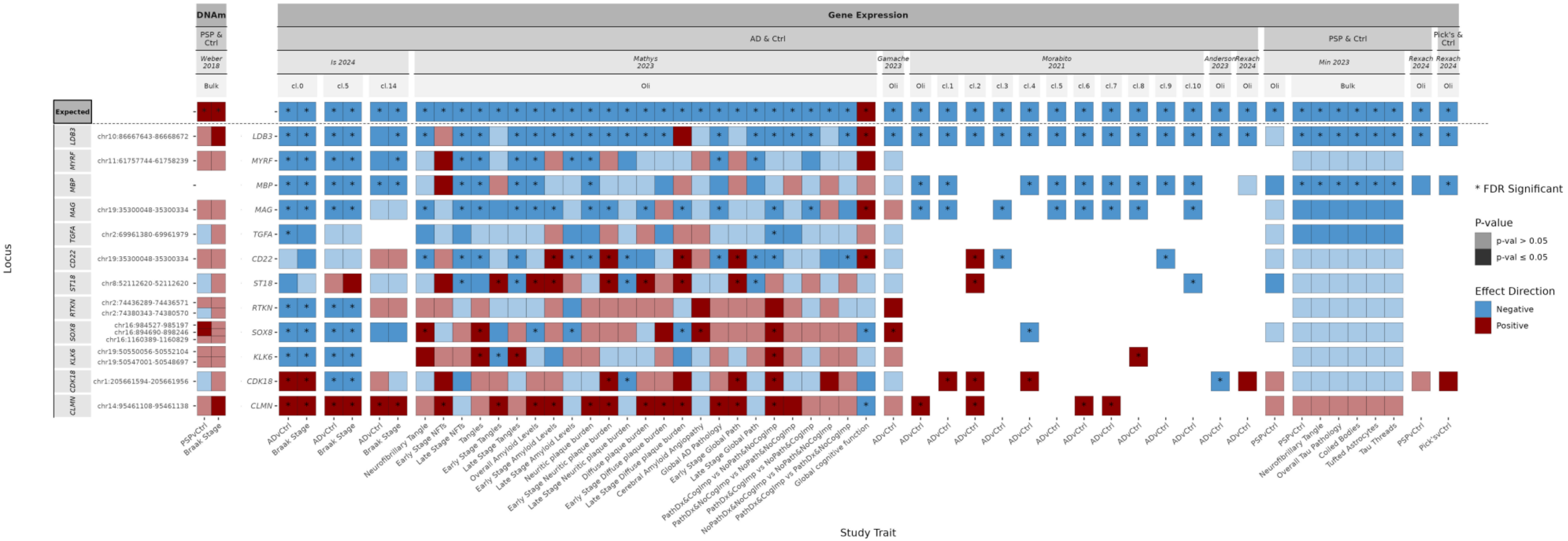

**Figure S10: Associations of rCpGm and genes of interest across multiple, previously published tau-related studies**.

Association results of top tau-related rCpGm or oligodendrocyte gene expression levels to multiple tau, amyloid, and cognition related endophenotypes pulled from multiple, recently published studies. Top X-axis is faceted in the first levels between whether DNAm or gene expression levels were tested, second the disease type of the study cohort examined, third the study examined, fourth whether bulk DNAm/RNAseq data, pseudobulk oligodendrocyte clusters from sn/scRNAseq data or specific oligodendrocyte assigned clusters were analyzed, and finally the bottom x-axis is the study trait investigated. The first row "Expected" depicts whether we expect a positive (red) or negative (blue) association based on our dataset. Locus annotations on the y-axis include the rCpGms defined by CpGm's present in the Weber *et al* 2018 study that overlap with our TCX rCpGms of interest. White areas indicate a result for that association was not reported from the study indicated.

**abs** = absolute value; **AD**= Alzheimer's Disease; **cl.** = cluster; **Ctrl** = Control; **DNAm** = DNA methylation; **ge** = Gene expression; **Oli** = Pseudobulk Oligodendrocytes; **PSP**= Progressive Supranuclear Palsy; **pTau** = phospho-tau; **p-val** = p-value; **TBS** = soluble tris buffered saline fraction; **tTau** = total tau; **TX**= membrane associated Triton-X fraction

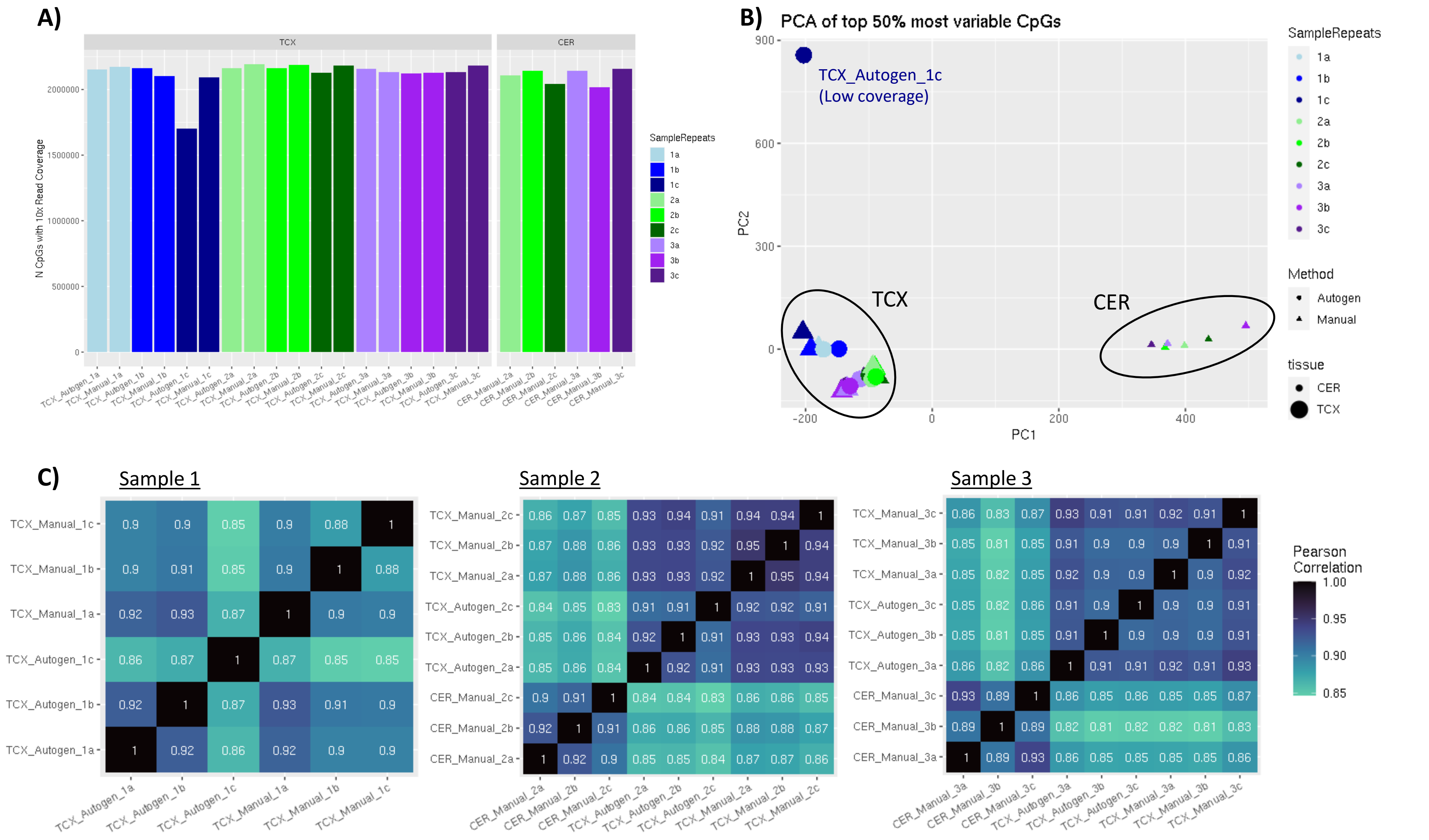


**Figure S11: Pilot experiment results comparing Autogen vs Manual DNA isolation methods on DNA methylation levels**. Samples were isolated using either the Autogen instrument or manually as detailed in the methods section in triplicate (a-c) for 3 samples (1-3). Libraries were prepared and quality controlled as detailed in the methods. A) Number of CpGs with 10x read coverage per sample included in the pilot experiment. Sample *TCX_Autogen_1c* has a much lower number of CpGs with 10x coverage indicating poor sample quality. B) PCA plot of the top 50% most variable CpG sites with 10x coverage. PC1 explains 10.9% of the variance while PC2 explains 6.8%. C) Person correlation of the top 50% most variable CpG sites with 10x coverage by sample.

**CER** = Cerebellum, **TCX** = Temporal Cortex
